# Transcription regulation by CarD in mycobacteria is guided by basal promoter kinetics

**DOI:** 10.1101/2023.03.16.533025

**Authors:** Dennis X. Zhu, Christina L. Stallings

## Abstract

Bacterial pathogens like *Mycobacterium tuberculosis* (*Mtb*) employ transcription factors to adapt their physiology to the diverse environments within their host. CarD is a conserved bacterial transcription factor that is essential for viability in *Mtb*. Unlike classical transcription factors that recognize promoters by binding to specific DNA sequence motifs, CarD binds directly to the RNA polymerase (RNAP) to stabilize the open complex intermediate (RP_o_) during transcription initiation. We previously showed using RNA-sequencing that CarD is capable of both activating and repressing transcription *in vivo*. However, it is unknown how CarD achieves promoter specific regulatory outcomes in *Mtb* despite binding indiscriminate of DNA sequence. We propose a model where CarD’s regulatory outcome depends on the promoter’s basal RP_o_ stability and test this model using *in vitro* transcription from a panel of promoters with varying levels of RP_o_ stability. We show that CarD directly activates full-length transcript production from the *Mtb* ribosomal RNA promoter *rrnA*P3 (AP3) and that the degree of transcription activation by CarD is negatively correlated with RP_o_ stability. Using targeted mutations in the extended −10 and discriminator region of AP3, we show that CarD directly represses transcription from promoters that form relatively stable RP_o_. DNA supercoiling also influenced RP_o_ stability and affected the direction of CarD regulation, indicating that the outcome of CarD activity can be regulated by factors beyond promoter sequence. Our results provide experimental evidence for how RNAP-binding transcription factors like CarD can exert specific regulatory outcomes based on the kinetic properties of a promoter.

## INTRODUCTION

Throughout their life cycle, bacteria must continuously adapt their physiology to respond to and survive in their changing environments. As such, the ability to sense environmental signals and transduce these cues into an appropriate physiological response is important for the virulence of pathogens such as *Mycobacterium tuberculosis* (*Mtb*), which face threats from both the host immune system and antibiotic treatment. Regulation of transcription initiation is a major mechanism by which bacteria adapt their gene expression in response to environmental stimuli. Transcription in bacteria is performed by a single RNA polymerase (RNAP) enzyme, which consists of a multi-subunit core enzyme that can bind to different sigma factors (σ) to form a holoenzyme and initiate promoter-specific transcription. *Mtb* devotes a significant fraction of its genome towards encoding numerous transcription factors that can regulate transcription initiation by altering the promoter specificity and recruitment of RNAP (1, 2). Classically, transcription factors are recruited to promoters by recognizing and binding a DNA sequence motif, which allows the factor to specifically regulate a subset of the genome. However, some bacteria also encode transcription factors that instead localize to promoter regions by binding directly to RNAP (3, 4). This class of transcription factors is best exemplified by the stringent response regulators DksA and guanosine (penta)tetraphosphate [(p)ppGpp], which bind to the *Escherichia coli* RNAP to directly activate or repress transcription from subsets of *E. coli* promoters (5). These factors exert promoter specific transcription regulation despite being unable to discriminate promoters at the level of binding. The prevailing hypothesis for the mechanism behind this promoter specificity postulates that these factors can potentiate different outcomes on transcription depending on the underlying initiation kinetics of a promoter (6). Recently, this hypothesis has also been applied to the regulatory mechanisms of other RNAP-binding transcription factors such as CarD (7, 8).

CarD is an RNAP-binding transcription regulator that is widely conserved across many eubacteria phyla and essential for viability in mycobacteria (9). CarD associates with transcription initiation complexes by binding directly to the RNAP β subunit through its N-terminal RNAP-interaction domain (RID) (9, 10). The CarD C-terminal DNA-binding domain (DBD) also interacts with DNA at the upstream fork of the transcription bubble in a sequence-independent manner (11–13). Numerous kinetic studies have demonstrated that CarD stabilizes the RNAP-promoter open complex (RP_o_) formed by the mycobacterial RNAP during transcription initiation (13–17). CarD accomplishes this through a two-tiered kinetic mechanism in which it binds to RNAP-promoter closed complexes (RP_c_) to increase the rate of DNA melting while also slowing the rate of bubble collapse (15). Furthermore, by stabilizing RP_o_, CarD slows the rate of promoter escape (18), which is a necessary step preceding full-length RNA synthesis. Due to its ability to stabilize RP_o_ *in vitro*, it was expected that CarD functioned generally as a transcription activator. However, although numerous studies have examined CarD’s effect on individual rate constants between transcription initiation intermediates (14, 15, 18), the composite effect of CarD’s kinetic mechanism on full-length RNA production remains unknown. Furthermore, while *in vitro* studies of CarD have utilized only a handful of promoters, primarily focusing on the *Mtb* ribosomal RNA promoter *rrnA*P3 (AP3) (12–16, 18–20), chromatin immunoprecipitation sequencing (ChIP-seq) in *Mycobacterium smegmatis* indicates that CarD co-localizes with the housekeeping sigma factor σ^A^ to promoter regions broadly across the mycobacterial genome (17, 21), leaving a gap in our understanding of CarD’s activity under different promoter contexts.

To characterize CarD’s role in transcription regulation throughout the mycobacterial genome, we previously performed RNA-sequencing (RNA-seq) on a set of *Mtb* strains expressing mutants of CarD that either impair or enhance its ability to stabilize RP_o_ *in vitro* (8). We discovered that altering CarD activity in *Mtb* led to both up-regulation and down-regulation of numerous protein-encoding transcripts, suggesting that CarD could function as either a transcriptional activator or a transcriptional repressor in different promoter contexts. Prior *in vitro* studies with *Rhodobacter sphaeroides* CarD and RNAP have shown that *Rsp*CarD activates transcription from promoters lacking a conserved T at the −7 position (22) and represses transcription from its own promoter (23). However, unlike Alphaproteobacteria like *R. sphaeroides*, which contain a T_-7_ at fewer than 50% of their promoters, most other bacterial phyla, including Actinobacteria like *Mtb*, have a T_-7_ at over 90% of their promoters (22), making it unlikely that the T_-7_ is a conserved mechanism of CarD promoter specificity. Instead, we previously proposed a model in which the outcome of CarD regulation is dependent on the basal transcription initiation kinetics at a given promoter (7, 8). Specifically, at unstable promoters that are rate-limited at the step of bubble opening, CarD would facilitate full-length RNA production by stabilizing RP_o_, while at stable promoters that are rate-limited at the step of promoter escape, CarD would make it more difficult for RNAP core enzyme to break contacts with promoter DNA. Herein, we directly test our model using *in vitro* transcription approaches to explore the relationship between RP_o_ stability and transcription regulation by CarD. We discover that both promoter DNA sequence and DNA topology influence the basal RP_o_ stability of a promoter and the regulatory outcome of CarD on transcription. In addition, we find that in the context of a promoter with high basal RP_o_ stability, CarD can directly repress transcription, marking the first demonstration of direct transcriptional repression by *Mtb* CarD. This work provides experimental evidence for how RNAP-binding transcription factors like CarD can potentiate multiple regulatory outcomes on transcription through a single kinetic mechanism.

## RESULTS

### CarD binding correlates with transcriptional regulation but not the direction of regulatory outcome

A fundamental feature in our model of CarD mechanism is that the regulatory outcome of CarD on a given mycobacterial promoter is determined based on differences in the basal transcription initiation kinetics of the promoter and not differences in CarD binding. This model is based on comparing ChIP-seq data from *M. smegmatis*, where CarD is present at almost all RNAP-σ^A^ transcription initiation complexes (17, 21), with RNA-seq data from *Mtb*, where mutation of CarD resulted in both up- and downregulation of gene expression (8). However, we cannot yet rule out the alternative hypothesis that CarD’s uniform localization pattern in *M. smegmatis* represents a unidirectional transcription activating mechanism for CarD in *M. smegmatis* in contrast to the bi-directional regulatory activity in *Mtb* that is suggested by our RNA-seq data.

To address this gap in our model, we performed an RNA-seq experiment in *M. smegmatis* that could be directly compared to the *M. smegmatis* ChIP-seq dataset. In our published *Mtb* RNA-seq experiment (8), we collected RNA from *Mtb* strains expressing mutant alleles of CarD with either weakened affinity for RNAP (CarD^R47E^), predicted weakened affinity for DNA (CarD^K125A^), or increased affinity for RNAP (CarD^I27F^ and CarD^I27W^). By collecting RNA from *Mtb* strains with mutations that target different domains of CarD, we were able to dissect how the respective interactions of CarD’s functional domains contributed to its role in regulating the *Mtb* transcriptome. To replicate this experimental design in *M. smegmatis*, we collected RNA from four strains of *M. smegmatis* with the native copy of *carD* deleted and expressing one of four different alleles of *Mtb* CarD: wild-type (WT) CarD (CarD^WT^), CarD^R25E^ (a RID mutant with weakened affinity for RNAP), CarD^K125E^ (a DBD mutant with weakened affinity for DNA), or CarD^I27W^ (a RID mutant with increased affinity to RNAP) as the only *carD* allele (**Table S1**). Similar to the CarD mutations used in our *Mtb* experiment, the CarD mutations that weaken its macromolecular interactions with RNAP or DNA (R25E and K125E) impair CarD’s ability to stabilize RP_o_ *in vitro* (13, 15), while the I27W mutation increases its affinity for RNAP and allows CarD to potentiate RP_o_-stabilization at lower concentrations (20). For each strain, we collected RNA from four biological replicates of exponentially growing cells in nutrient replete conditions for sequencing. Two replicates (CarD^R25E^-1 and CarD^K125E^-4) were identified as outliers following principal component analysis (PCA) and were discarded from downstream analysis (**Fig. S1**).

In all three strains with mutations in *carD,* over 25% of the 6,716 coding genes in *M. smegmatis* Mc^2^155 were significantly differentially expressed (*P_adj_* < 0.05) in comparison to the CarD^WT^ strain (**Fig. 1A**, **Table S1**). The number of differentially expressed genes in the CarD^R25E^ (2909 genes) and CarD^K125E^ (2901 genes) *M. smegmatis* strains is similar to the number of differentially expressed genes in the CarD^R47E^ (2877 genes) and CarD^K125A^ (2690 genes) *Mtb* strains (8). However, homologous genes between the two species showed little correlation in their transcript expression patterns (**Fig. S2, Table S2**), suggesting that CarD does not simply regulate a subset of homologous genes conserved between *Mtb* and *M. smegmatis*. Each of the *M. smegmatis* CarD mutant strains exhibited a similar number of up-regulated genes as down-regulated genes (**Fig. 1A**), following the same pattern as the *Mtb* CarD mutant strains (8) and suggesting that CarD is capable of potentiating both transcriptional activation and repression in *M. smegmatis*. Importantly, the strains did not show significant differences in the total amount of RNA per cell (**Fig. 1B**), suggesting that the transcript abundance differences measured in the CarD mutant strains represent local changes in transcription at specific genes rather than a global decrease in RNA production within the cell that would be expected if CarD functioned strictly as a transcriptional activator.

**Figure 1.**
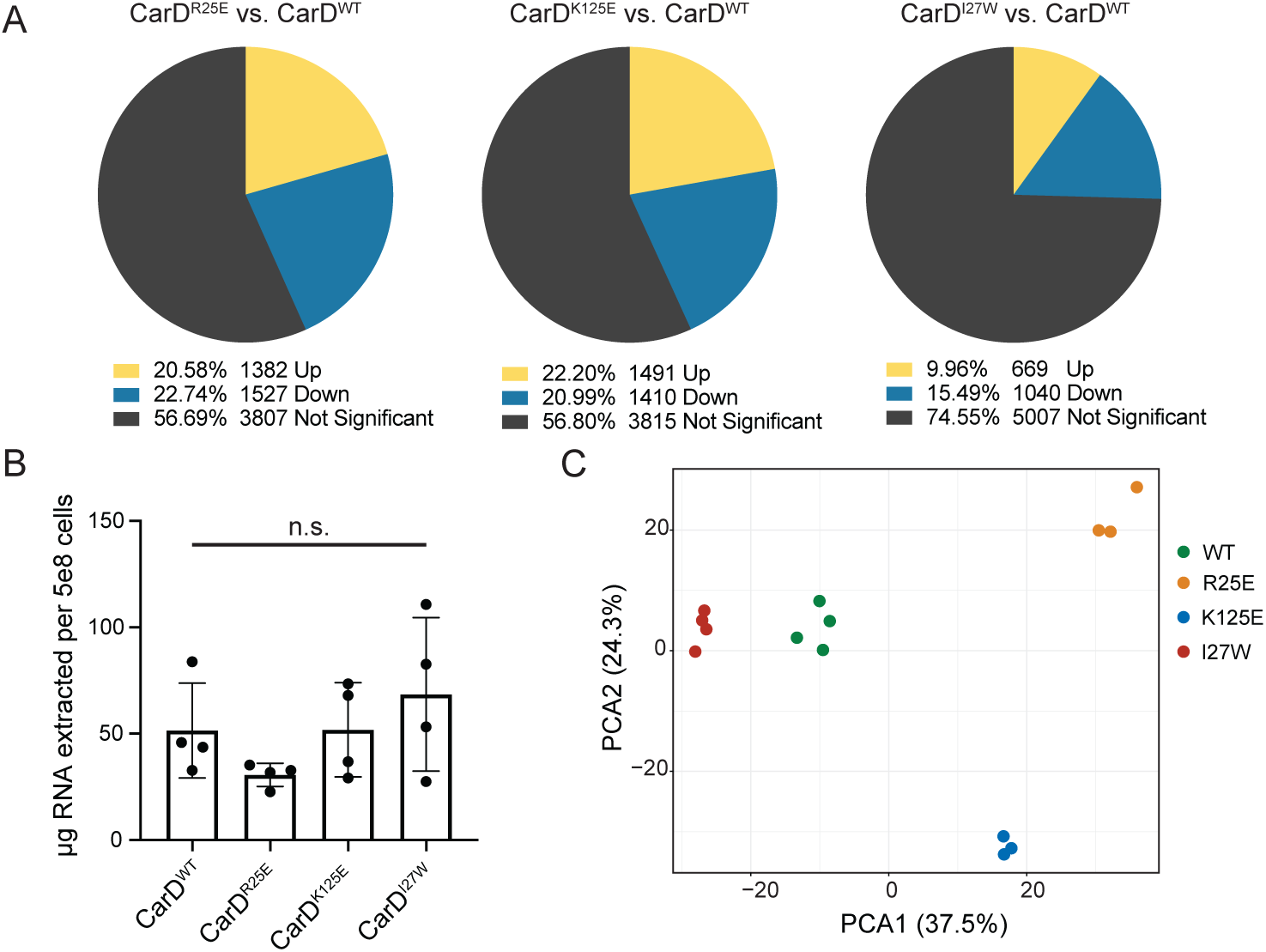
*M. smegmatis* strains encoding point mutants of CarD display broad changes in transcript expression. (A) Pie charts displaying the percentage of *M. smegmatis* coding genes that were significantly differentially expressed (*p_adj_* < 0.05) in each CarD mutant strain relative to CarD^WT^. (B) RNA content for *M. smegmatis* strains expressing different alleles of CarD calculated from total RNA weight harvested from four biological replicates divided by estimated number of cells collected. Each bar represents the mean ± standard deviation (SD). Group means were compared using a one-way ANOVA and determined to be not significantly different (*p* = 0.228). (C) Principal component analysis of RNA sequencing samples based on read counts of 6,716 *M. smegmatis* MC^2^155 coding genes. The first two principal components (PC1 and PC2), which account for 37.5% and 24.3% of the variance, respectively, define the x- and y-axis, respectively.

The transcriptomic relationship between different CarD mutant strains in *M. smegmatis* was also consistent with the relationships we observed in our *Mtb* dataset (8). PCA of the RNAseq data illustrated that the *M. smegmatis* sample replicates clustered tightly with each other based on CarD genotype and samples from CarD mutant strains with impaired RP_o_-stabilizing activity (CarD^R25E^ and CarD^K125E^) separated from the strain with enhanced RP_o_-stabilizing activity (CarD^I27W^) along the first principal component (**Fig. 1C**), demonstrating consistency between replicates from the same genotype and suggesting that altered CarD RP_o_-stabilizing activity contributes to transcript abundance changes in the mutant bacteria. In addition, the CarD^R25E^ and CarD^I27W^ strains, which encode CarD RID mutants with impaired or enhanced RP_o_-stabilization *in vitro*, respectively, displayed largely opposite transcriptomic changes (**Fig. S3A**), similar to the RID mutants in *Mtb* (8). In the PCA, the CarD^K125E^ samples separated from all other samples along the second principal component (**Fig. 1A**) and the direction of transcript abundance changes in the DBD mutant CarD^K125E^ samples correlated poorly with the transcript abundance changes in the RID mutant CarD^R25E^ samples (R^2^ = 0.351) (**Fig. S3B-C**). This is in contrast to the tight correlation between CarD RID and DBD mutants in *Mtb* (8) and may suggest that mutations in the DBD and RID have unique effects on CarD’s regulatory function in *M. smegmatis*.

The ChIP-seq dataset shows that CarD is present when RNAP-σ^A^ is also found, supporting a model that CarD is present at the promoters of both up- and downregulated genes. However, our data is also compatible with an alternative model in which CarD acts directly as a monotonic transcriptional activator and genes that appear to be transcriptionally “repressed” by CarD are expressed at lower levels in WT bacteria due to decreased RNAP occupancy at non-CarD-activated promoters. If this alternative model were true, then we would expect to find RNAP-σ^A^/CarD binding sites overlapping with transcription start sites (TSSs) ascribed to the transcriptionally “activated” promoters but absent from TSSs ascribed to transcriptionally “repressed” promoters. To examine the overlap between CarD binding sites and CarD-regulated transcripts, we used our RNA-seq dataset to identify a list of *M. smegmatis* genes whose transcript abundance was likely directly responsive to altered CarD-mediated RP_o_ stabilization activity based on having opposite expression patterns in CarD^R25E^ versus CarD^I27W^. To avoid internal genes within operons, we focused our analysis on 2,917 *M. smegmatis* genes directly downstream of a primary TSS (24) and categorized them into one of four classes (**Table S3**). TSSs associated with genes that were significantly down-regulated (*P_adj_* < 0.05) in CarD^R25E^ and significantly up-regulated in CarD^I27W^ were classified as ‘Activated’ by CarD (n=117) while TSSs associated with genes that were significantly up-regulated in CarD^R25E^ and significantly down-regulated in CarD^I27W^ were classified as ‘Repressed’ by CarD (n=153). TSSs associated with genes that were significantly differentially expressed in both CarD^R25E^ and CarD^I27W^ but in the same direction relative to wild-type were classified as ‘Uncategorized’ (n=222) because their expression profile does not reflect the divergent expression pattern expected between CarD mutants with opposing effects on RP_o_-stabilization *in vitro*. Lastly, any TSSs that were not significantly differentially expressed in both CarD^R25E^ and CarD^I27W^ were categorized as ‘Not Significant’ (n=2425). We re-analyzed our previous ChIP-seq dataset (17, 21) and identified 1857 unique CarD binding sites across two biological replicates (**Table S3**). To avoid broad binding regions that may represent multiple, overlapping CarD binding sites, we focused on 1796 CarD binding sites less than or equal to 1000 base pairs (bp) in width. Of these 1796 CarD binding sites, 1390 sites (77.4%) overlapped with at least one mapped TSS in *M. smegmatis* and 1129 sites (62.8%) overlapped with a primary TSS associated with a protein-encoding gene (24). We examined the overlap between CarD binding sites and TSSs that were significantly differentially expressed in both CarD^R25E^ and CarD^I27W^ and found that 57.9% (285/492) of these TSSs were associated with CarD binding (**Table 1**). Among the differentially expressed genes, 53.0% (62/117) of ‘Activated’ TSSs and 67.3% (103/153) of ‘Repressed’ TSSs overlapped with a CarD binding site (**Table 1**). Thus, CarD binding is associated with transcriptional regulation of *M. smegmatis* promoters *in vivo* but is not correlated with the direction of regulation. A similar analysis was performed in the ɑ-proteobacterium *Caulobacter crescentus* to identify the direct regulon of CdnL (the *C. crescentus* homolog of CarD) (25). Like our results, CdnL localized to promoter regions of both genes that were up-regulated and genes that were down-regulated in a Δ*cdnL* strain, but a vast majority of differentially expressed genes were not associated with CdnL binding, suggesting a broader effect of indirect regulation in *C. crescentus*. Together, these data support the model that CarD is broadly localized to mycobacterial promoters through its interaction with RNAP but that the regulatory outcome of CarD activity is not determined by occupancy.

**Table 1.** CarD binding is associated with both activated and repressed transcription start sites (TSSs). ‘Differentially Expressed’ TSSs are those TSSs associated with genes that were significantly differentially expressed (p*_adj_*<0.05) in both CarD^R25E^ and CarD^I27W^ relative to CarD^WT^. TSSs associated with genes that were not significantly differentially expressed in both mutant strains were categorized as ‘Not Significant’. P-values from a hypergeometric test are listed. Differentially expressed genes were categorized as: ‘Activated’ if it was down-regulated in CarD^R25E^ and up-regulated in CarD^I27W^, ‘Repressed’ if it was up-regulated in CarD^R25E^ and down-regulated in CarD^I27W^, or ‘Uncategorized’ if it was differentially expressed in the same direction in both mutant strains.

|  | Differentially Expressed | Not Significant | Activated | Repressed | Uncategorized | TOTAL |
| --- | --- | --- | --- | --- | --- | --- |
| Overlapping with CarD binding site | 285<br>(57.9%) | 1089<br>(44.9%) | 62<br>(53.0%) | 103<br>(67.3%) | 120<br>(54.0%) | 1374<br>(47.1%) |
| Total # TSSs | 492 | 2425 | 117 | 153 | 222 | 2917 |

### CarD directly activates transcription from the *Mtb* ribosomal RNA promoter *rrnA*P3

To test our model that the outcome of CarD’s RP_o_ stabilizing activity on transcript production depends on the basal promoter kinetics, we used *in vitro* transcription methods to measure the direct effects of CarD on transcript production. Although several studies have proposed that CarD activates transcription from the *Mtb* AP3 promoter based on *in vitro* three-nucleotide transcription assays (16, 20) and real-time fluorescence assays (14, 15) that report RP_o_ lifetime, full-length transcript production has never been directly measured. To assess full-length RNA production, we performed multi-round *in vitro* transcription assays by incubating recombinantly purified *Mtb*RNAP-σ^A^ holoenzyme with a linear DNA fragment containing the *Mtb* AP3 promoter (from −39 to +4 with respect to the TSS) driving transcription of a 164 nucleotide RNA product. The addition of a saturating concentration of WT CarD (25:1 molar ratio CarD:RNAP (15)) activated transcription from the AP3 promoter ∼8-fold compared to reactions with no factor added (**Fig. 2A**). To investigate how CarD’s RP_o_-stabilizing activity relates to transcriptional activation, we repeated the multi-round *in vitro* transcription assays with CarD mutants impaired in their ability to stabilize RP_o_ *in vitro* (CarD^R25E^, CarD^R47E^, CarD^K125A^, and CarD^K125E^) (13, 15). All four of the CarD mutants activated transcription from AP3 compared to reactions with no factor, but the degree of activation by each mutant was reduced compared to WT CarD (**Fig. 2A**), suggesting that CarD’s RP_o_-stabilizing activity underlies its ability to activate transcription from AP3. In addition, the degree to which each CarD mutant attenuated transcript production correlated with how severe the impact was on the CarD macromolecular interactions with RNAP (10) or DNA (13). In contrast, CarD^I27W^, which has increased affinity for RNAP and is able to stabilize RP_o_ at lower concentrations than CarD^WT^ (20), activated transcription from AP3 to a greater degree than CarD^WT^ at concentrations below where CarD^WT^ is saturating (5:1 molar ratio CarD:RNAP) (**Fig. 2B**), further demonstrating the association between CarD’s RP_o_-stabilizing activity and activation of transcript production. Collectively, these results demonstrate that CarD activates full-length RNA production *in vitro* from AP3 and this transcription activation is dependent on the RP_o_-stabilizing activity of CarD.

**Figure 2.**
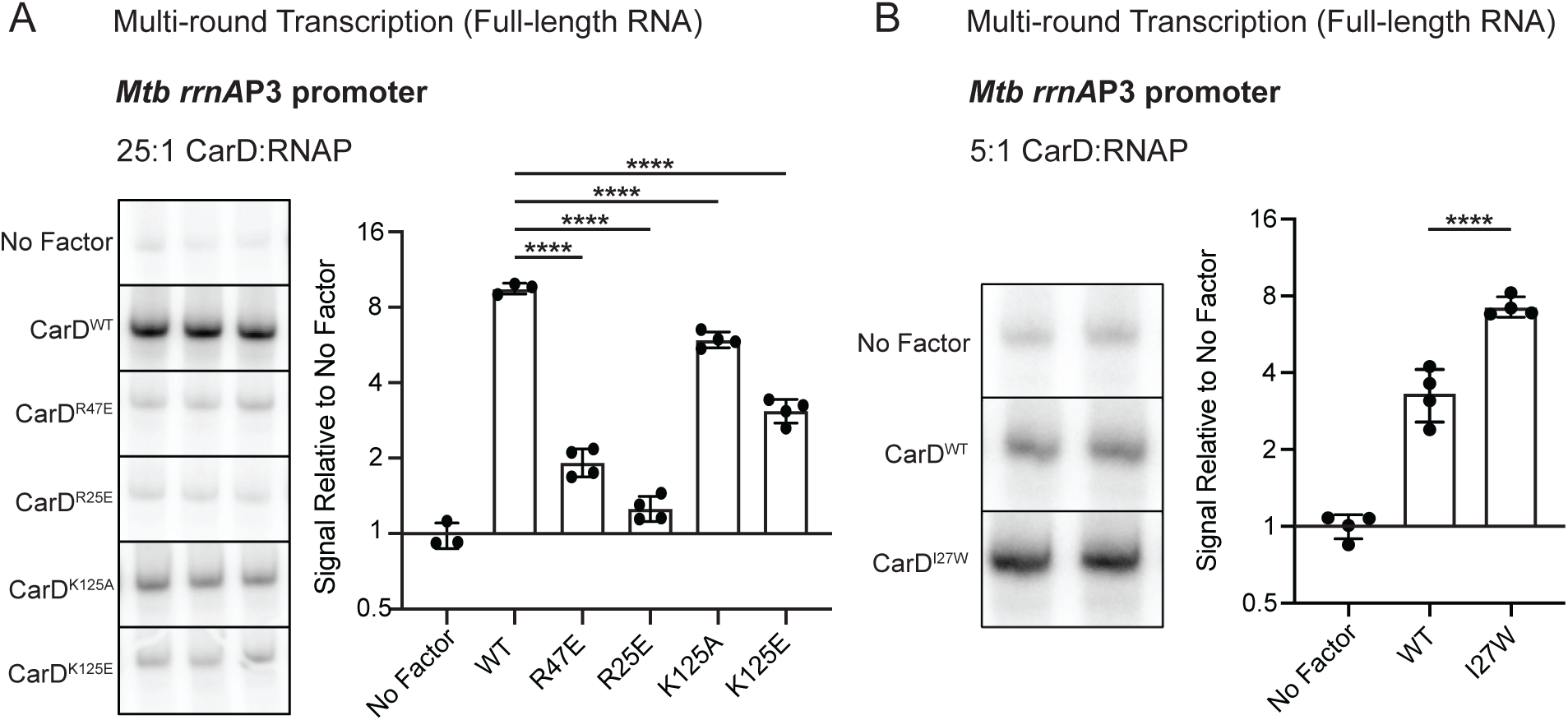
CarD activates transcription from the *Mtb* ribosomal RNA promoter AP3, and mutations to either the RNA polymerase (RNAP) interaction domain (RID) or DNA-binding domain (DBD) impair this activity *in vitro*. (A) Representative gel images from multi-round *in vitro* transcription reactions of *Mtb*RNAP-σ^A^ holoenzyme on linear DNA templates encoding AP3 with either no factor, wild-type CarD (CarD^WT^), one of two RID mutants (CarD^R47E^ or CarD^R25E^), or one of two DBD mutants (CarD^K125A^ or CarD^K125E^). In all reactions with factor, CarD is added at a 25:1 molar ratio to RNAP holoenzyme. The bar graph displays the mean transcript signal intensity relative to ‘No Factor’ ± standard deviation (SD). N=3-4 independent reactions for each condition. (B) Representative gel images from multi-round *in vitro* transcription reactions of *Mtb*RNAP-σ^A^ holoenzyme on AP3 with either no factor, CarD^WT^, or a RID mutant with higher affinity for RNAP (CarD^I27W^). In all reactions with factor, CarD is added at a sub-saturating concentration of 5:1 molar ratio to RNAP holoenzyme. The bar graph displays the mean transcript signal intensity relative to ‘No Factor’ ± SD. N=4 independent reactions for each condition. (A and B) Mean fold-change values were compared using a one-way ANOVA followed by post-hoc Dunnett’s tests comparing the mean of each mutant CarD allele to CarD^WT^; **** = *p*<0.0001. The full ANOVA results are listed in **Table S4**.

### Additional promoter DNA-RNAP interactions increase basal RP_o_ stability and push CarD towards transcriptional repression

To test the hypothesis that the degree of transcriptional activation by CarD is inversely correlated with the basal RP_o_ stability of a promoter, we explored CarD’s direct regulatory effect on transcription from a set of promoters with varying levels of basal RP_o_ stability. Transcription initiation kinetics and RP_o_ lifetime are highly dependent on promoter DNA sequence (26). In the RP_o_ intermediate, the promoter DNA makes multiple sequence-specific contacts with regions of the RNAP holoenzyme to stabilize the transcription bubble (26–28). The *Mtb* RNAP-σ^A^ holoenzyme and WT AP3 (AP3_WT_) promoter form a relatively unstable RP_o_ (15, 16) that is stabilized by CarD to lead to activation of transcription *in vitro* **(Fig. 2)**. We, therefore, used the AP3 promoter sequence as a starting point to generate four additional promoter templates (AP3_EcoExt_, AP3_MycoExt_, AP3_Discr_, and AP3_Stable_) with higher levels of basal RP_o_ stability by making targeted sequence mutations that would add or optimize predicted DNA-RNAP interactions in RP_o_ (**Fig. 3A**). AP3_WT_ contains near consensus sequence motifs in the −35 and −10 elements (29), which are highly conserved promoter elements that interact with σ region 4 and 2, respectively (30–32), so we did not target these regions in our study. In AP3_EcoExt_, we mutated the base at position −14 to a G to introduce a T_-15_G_-14_ motif that represents an extended −10 element that was first identified in *E. coli* (33). In addition to the classical *E. coli*-like extended −10 motif, many mycobacterial promoters instead contain a G at position −13 that is associated with promoter strength and RP_o_ formation in DNase I footprinting studies (34). Thus, we also generated AP3_MycoExt_, which is mutated to include a G_-13_ upstream of the −10 element. Both G_-14_ and G_-13_ are positioned to interact with a conserved glutamic acid residue in σ^A^ region 3.0 in the mycobacterial RP_o_ (14, 34, 35). AP3_Discr_ is mutated to introduce a G_-6_GGA_-3_ motif in the discriminator region immediately downstream of the −10 hexamer that allows for optimal binding with σ^A^ region 1.2 (36–38). AP3_Stable_ is mutated to include the mutations made in AP3_EcoExt_ and AP3_Discr_ as well as a deletion of a T at position −17 to reduce the length of the spacer region between the −35 and −10 hexamers from 18-bp in AP3_WT_ to 17-bp. A spacer length of 17-bp allows for optimal interactions of the −35 and −10 hexamers with σ^A^ (39).

**Figure 3.**
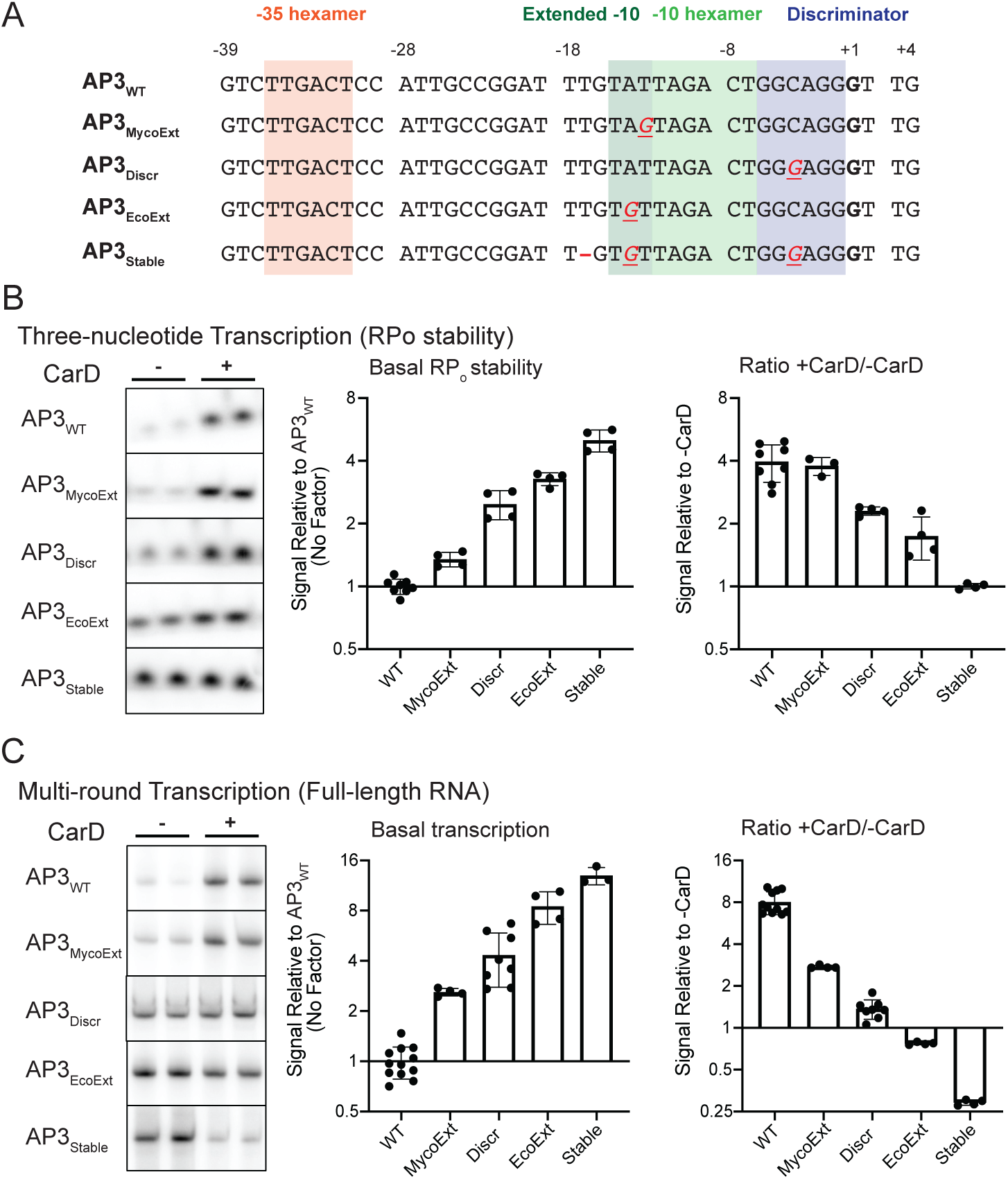
Promoter sequences that introduce additional interactions between promoter DNA and RNAP in the open complex increase basal RP_o_ stability and shift the regulatory outcome of CarD towards transcriptional repression. (A) Promoter sequences of the wild-type *Mtb rrnA*P3 promoter (AP3_WT_) and four variants with sequence mutations that add predicted interactions between promoter DNA and RNAP in RP_o_. Sequences from the −39 to +4 position relative to the transcription start site (+1, bolded) are shown. In the non-WT sequences, DNA bases that are altered from the WT sequence are underlined and colored red. A “—” indicates that a base was deleted. (B) Representative gels showing [^32^P]-labeled three-nucleotide transcription products formed by *Mtb*RNAP-σ^A^ from linear DNA templates encoding AP3_WT_ or one of the four AP3 variants either in the absence or presence of CarD. (C) Representative gels from multiround transcription assays showing 164 nucleotide [^32^P]-labeled RNA transcripts produced by *Mtb*RNAP-σ^A^ from linear DNA templates encoding AP3_WT_ or one of the four AP3 variants either in the absence or presence of CarD. (B and C) Bar graphs display (left) the mean basal signal intensity relative to AP3_WT_ ± standard deviation (SD) and (right) the mean ratio of signal intensity +CarD/-CarD for each promoter ± SD Group means were compared by one-way ANOVA *p*<0.0001. The full results of pairwise comparisons are listed in **Table S4**.

To measure the basal RP_o_ stability of RNAP-σ^A^ and the AP3 promoter variants, we performed *in vitro* three-nucleotide transcription initiation assays (16, 20) in the absence of CarD by incubating *Mtb*RNAP-σ^A^ holoenzyme with linear promoter DNA fragments in the presence of a GpU dinucleotide and UTP. In these reactions, the RNAP-σ^A^ holoenzyme can synthesize a three nucleotide ‘GUU’ RNA transcript but cannot undergo promoter escape, allowing us to assess relative RP_o_ lifetimes by using the amount of three nucleotide product as a proxy. We found that all the promoter variants with additional predicted DNA-RNAP contacts exhibited higher basal levels of RP_o_ stability compared to AP3_WT_ (**Fig. 3B**). The most stable variant AP3_Stable_ displayed 8-fold higher basal RP_o_ stability relative to AP3_WT_. To quantify the effect of CarD on RP_o_ stability from these promoter variants, we also performed three-nucleotide transcription assays in the presence of WT CarD protein (**Fig. 3B**). On AP3_WT_, CarD increased the amount of three nucelotide product by roughly 4-fold over reactions with no factor. As the basal RP_o_ stability of promoter variants increased, the degree of RP_o_ stabilization by CarD decreased to the point that on AP3_Stable_ the addition of CarD resulted in no detectable difference in the amount of three nucleotide product.

Having established a set of promoters with different basal RP_o_ stability levels that range over nearly one order of magnitude, we performed multi-round *in vitro* transcription reactions using these AP3 promoter variants in the presence or absence of CarD to investigate the relationship between the basal RP_o_ stability of a promoter and transcriptional regulation by CarD (**Fig. 3C**). We discovered that across the AP3 promoter variants, basal RP_o_ stability positively correlated with full length transcript production in the absence of CarD but negatively correlated with transcriptional activation by CarD. Indeed, the two promoters with the highest levels of basal RP_o_ stability (AP3_EcoExt_ and AP3_Stable_) were transcriptionally repressed by CarD, consistent with the predictions of our model and providing the first *in vitro* evidence of direct transcription repression by *Mtb* CarD.

### Basal RP_o_ stability and CarD regulatory outcome are influenced by discriminator region guanosine + cytosine base pair frequency

In addition to forming direct interactions with the polymerase, promoter DNA sequences can also influence RP_o_ stability by affecting the chemical properties of the DNA molecule. For example, guanosine + cytosine base pairs in the discriminator region impose a kinetic barrier to DNA untwisting and unwinding during the formation of the transcription bubble due to their greater base-pairing and base-stacking stability compared adenosine + thymine base pairs (40, 41). Discriminator guanosine + cytosine base pair frequency (G+C%) is inversely correlated with RP_o_ stability (42) and has been shown to be a determinant of transcription control by DksA/(p)ppGpp (43, 44). To determine if changing the RP_o_ stability by modifying the G+C% of the discriminator affects the outcome of CarD activity on transcript production, we generated a set of AP3 promoter variants (AP3_Discr1_ – AP3_Discr5_) in which the discriminator region G+C% is titrated from 100% (AP3_Discr1_) to 16.7% (AP3_Discr5_) (**Fig. 4A**). We observed a negative correlation between discriminator G+C% and basal RP_o_ stability as measured by three-nucleotide transcription assays (**Fig. 4B**). CarD increased three nucleotide RNA production from all promoter variants tested, but the magnitude of RP_o_ stabilization by CarD displayed a negative correlation with basal RP_o_ stability across AP3 variants as the discriminator G+C% was titrated (**Fig. 4B**).

**Figure 4.**
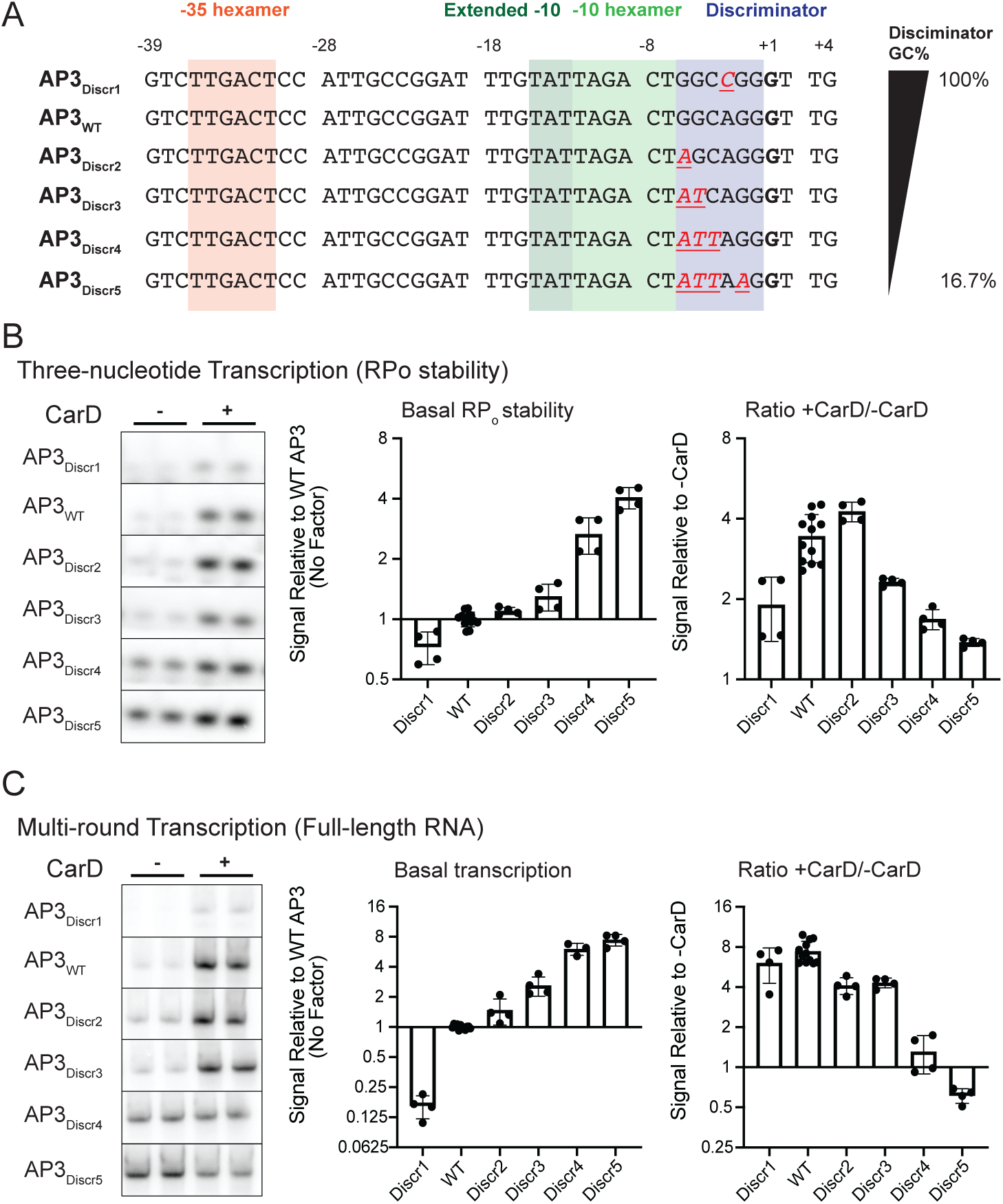
Discriminator GC% negatively correlates with basal RP_o_ stability and influences CarD regulatory outcome. (A) Promoter sequences of the wild-type *Mtb rrnA*P3 promoter (AP3_WT_) and five variants with sequence mutations that either increase or decrease the percentage of G or C bases in the discriminator. Sequences from the - 39 to +4 position relative to the transcription start site (+1, bolded) are shown. In the non-WT sequences, DNA bases that are altered from the WT sequence are underlined and colored red. (B) Representative gels showing [^32^P]-labeled three-nucleotide transcription products formed by *Mtb*RNAP-σ^A^ from linear DNA templates encoding AP3_WT_ or one of the four AP3 variants either in the absence or presence of CarD. (C) Representative gels from multi-round transcription assays showing 164 nucleotide [^32^P]-labeled RNA transcripts produced by *Mtb*RNAP-σ^A^ from linear DNA templates encoding AP3_WT_ or one of the four AP3 variants either in the absence or presence of CarD. (B and C) Bar graphs display (left) the mean basal signal intensity (in the absence of CarD) relative to AP3_WT_ ± standard deviation (SD) and (right) the mean ratio of signal intensity +CarD/-CarD for each promoter ± SD Group means were compared by one-way ANOVA *p*<0.0001. The full results of pairwise comparisons are listed in **Table S4**.

Discriminator G+C% of the AP3 variants was also negatively correlated with basal transcript production in the multi-round transcription assay and the magnitude of transcription activation by CarD decreased as discriminator G+C% decreased (**Fig. 4C**). On the AP3 variant with the lowest discriminator G+C% and highest basal RP_o_ stability (AP3_Discr5_), CarD decreased transcript production, further supporting that promoters with high basal RP_o_ stability can be transcriptionally repressed by CarD. Collectively, our experiments show that promoter sequence motifs that increase basal RP_o_ stability decrease the magnitude of transcriptional activation by CarD and can lead to transcriptional repression in the most stable RP_o_ contexts.

### Promoter sequences that form more stable RP_o_ are associated with transcription repression by CarD *in vitro* and *in vivo*

We show that base substitutions in the spacer region, extended −10 region (**Fig. 3**), and discriminator (**Fig. 4**) can affect full-length transcript production and the direction of CarD regulation. To directly examine whether differences in relative RP_o_ stability could explain the outcomes in transcript production and CarD regulation we performed a linear regression analysis across all of our promoter templates (**Fig. 5**). For this analysis, the relative RP_o_ stability and relative transcription strength of each promoter variant was normalized to AP3_WT_. In the absence of CarD, the rate of full-length transcript production shows a roughly linear positive correlation with the relative RP_o_ stability (**Fig. 5A**). In contrast, the log_2_ ratio of transcript production in multi-round transcription reactions +/- CarD shows a roughly linear inverse correlation with increasing RP_o_ stability, with the most stable promoter variants (AP3_EcoExt_, AP3_Stable_, and AP3_Discr5_) being transcriptionally repressed by CarD (**Fig. 5B**). The robust relationship across multiple promoter variants suggests that RP_o_ stability is a fundamental determinant of full-length transcript production and CarD regulatory outcome. Collectively, our experiments illustrate a relationship between RP_o_ stability, transcription strength, and CarD regulation and demonstrate that transcription factors like CarD can discriminate promoters based on their basal kinetic features to potentiate bidirectional outcomes in transcription regulation via a single kinetic mechanism.

**Figure 5.**
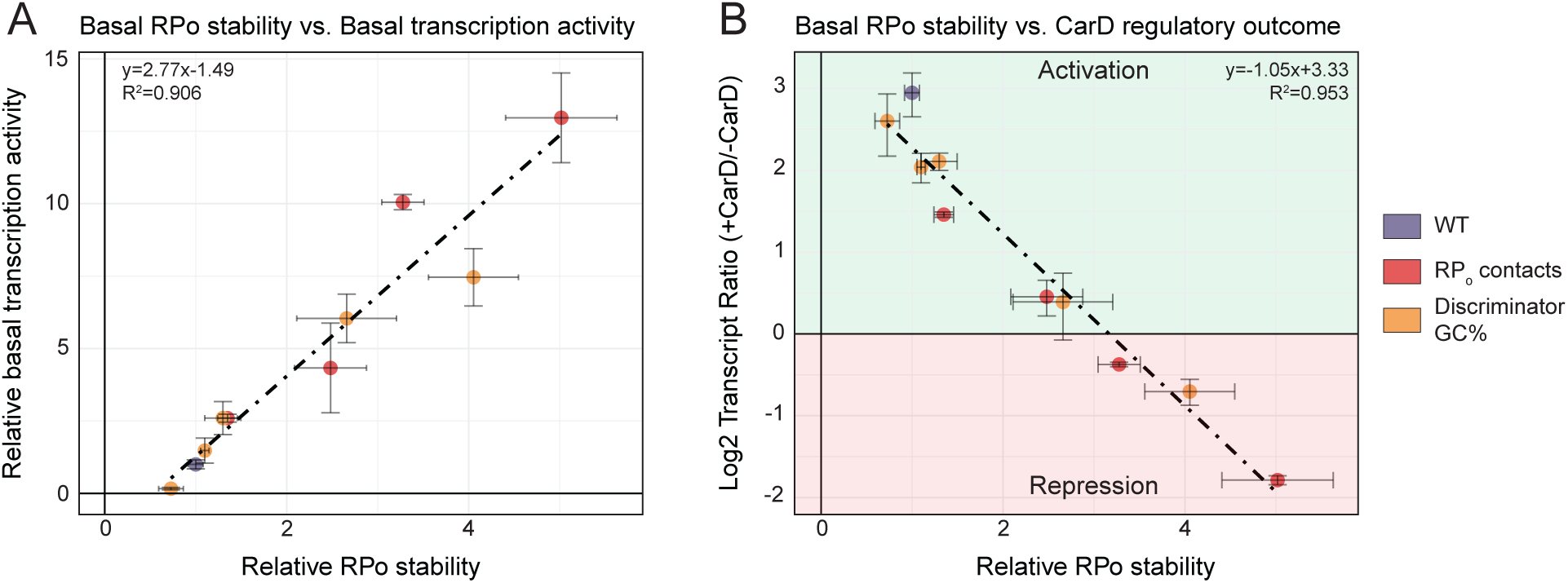
The basal RP_o_ stability of a promoter is positively correlated with basal transcription activity but negatively correlated with transcription activation by CarD. (A) Dot plot showing the relationship between the basal RP_o_ stability of AP3 promoter variants relative to AP3_WT_ on the x-axis versus basal transcription activity relative to AP3_WT_ on the y-axis. Each point represents a variant of the AP3 promoter and is colored based on whether it represents the WT promoter, a sequence with mutations that affect RNAP-DNA interactions in RP_o_ (RP_o_ contacts), or a sequence with mutations that affect the discriminator region GC% (Discriminator GC%). The position of each point represents the mean values from at least N=4 experiments and the error bars represent standard deviations. The dashed line and text represent the results of a linear regression analysis. (B) Dot plot showing the relationship between the basal RP_o_ stability of AP3 promoter variants versus the log_2_ ratio of transcript production in reactions ±CarD on the y-axis. Positive ‘Log_2_ Transcript Ratio’ values indicate transcription activation while negative values indicate transcriptional repression.

Through our *in vitro* transcription experiments, we have identified multiple promoter sequence motifs associated with high RP_o_ stability *in vitro*. If our model is generally applicable to transcription from mycobacterial promoters throughout the genome, then we would expect to find an association between DNA sequence motifs associated with RP_o_ stability and transcriptional repression by CarD. To interrogate this prediction, we examined the prevalence of a consensus extended −10 motif (T_-15_G_-14_N_-13_) and discriminator GC% in promoters that were differentially expressed in *our Mtb* and *M. smegmatis* RNA-seq datasets (**Table S5**). Since all of our *in vitro* experiments were performed in the context of a *Mtb*RNAP-σ^A^ holoenzyme, we limited our bioinformatic analysis to promoters containing a A_-11_NNNT_-7_ motif representing the consensus σ^A^ −10 element (24, 29, 45, 46), which comprised 90.5% (1609/1778) and 82.5% (2511/3043) of the primary TSSs in *Mtb* (45) and *M. smegmatis* (24), respectively. Indeed, in *Mtb*, promoters that were predicted to be repressed by CarD based on our RNA-seq data were over-enriched for extended −10 elements while promoters predicted to be activated by CarD were under-enriched for extended −10 elements relative to the genome-wide proportion of this feature (**Fig. S4A**). A similar trend was true of the proportion of promoters containing extended −10 elements in *M. smegmatis*, but the difference in proportions between CarD regulated promoters and the genome-wide distribution was not statistically significant (**Fig. S4B**). In both species, promoters that were predicted to be repressed by CarD contained significantly more GC-rich discriminator regions than promoters predicted to be activated by CarD (**Fig. S4C-D**). The association of stable RP_o_ DNA sequence signatures with genes that are inferred to be repressed by CarD *in vivo* support that the regulatory mechanisms that we demonstrate *in vitro* could be relevant to gene expression *in vivo*.

### DNA topology can influence the regulatory outcome of CarD activity

In mycobacteria, CarD transcript levels increase in response to double-stranded DNA breaks and genotoxic stress (9), suggesting that the dynamics of CarD regulation may be important for responding to these environmental cues. DNA breaks in the chromosome can relieve local regions of DNA supercoiling. The supercoiling state of promoters is tightly connected to transcriptional activity *in vivo*, as positive or negative supercoiling can inhibit or enhance RP_o_ formation, respectively (47, 48). Thus, we sought to test the relationship between promoter topology and CarD regulation. We generated a set of templates with identical DNA sequence but varied molecular topology by cloning the AP3_WT_ promoter into a negatively supercoiled plasmid and incubating the plasmid with either a single-cutting endonuclease to produce a linear “cut” DNA molecule, a nicking endonuclease to produce a circular “nicked” DNA molecular, or with no enzyme to maintain a supercoiled control (mock treated) (**Fig. 6A**). We performed *in vitro* three-nucleotide transcription assays using the topologically distinct DNA templates and found that negative supercoiling contributes to a ∼6-fold increase in basal RP_o_ stability compared to a linear “cut” DNA template containing the same promoter sequence (**Fig. 6B**). In addition, the “nicked” DNA template exhibited a similar basal RP_o_ stability to the “cut” DNA template, indicating that the higher RP_o_ stability observed in the “mock” template is a result of supercoiling and not the circular shape of the molecule. The addition of CarD decreased the amount of three nucleotide transcript produced with the supercoiled “mock” DNA template. This result could indicate that CarD inhibits progression from RP_o_ towards an initial transcribing complex intermediate (RP_itc_) that synthesizes the three nucleotide product quantified in these assays (18). The basal RP_o_ stabilities of the “cut”, “nicked”, and “mock” AP3_WT_ DNA templates correlated with the basal transcriptional activity of the promoter, where promoter templates with high basal RP_o_ stability also showed high levels of basal transcript production (**Fig. 6C**). Furthermore, CarD activated transcription from the “cut” and “nicked” DNA templates but repressed transcription from the supercoiled “mock” DNA template, which has a higher basal RP_o_ stability relative to the “cut” and “nicked” molecules. These data demonstrate a single promoter DNA sequence can exhibit varying levels of basal RP_o_ stability based on DNA supercoiling, and this supercoiling-dependent change in RP_o_ stability can change the regulatory outcome of CarD on transcription. While the DNA sequence of a given promoter is constant within the genome, the topology of the DNA molecule can change over the lifetime of the cell. Thus, our findings reveal an additional layer of complexity in CarD’s regulatory mechanism and could help explain how CarD expression *in vivo* could lead to differential gene expression outcomes in different conditions.

**Figure 6.**
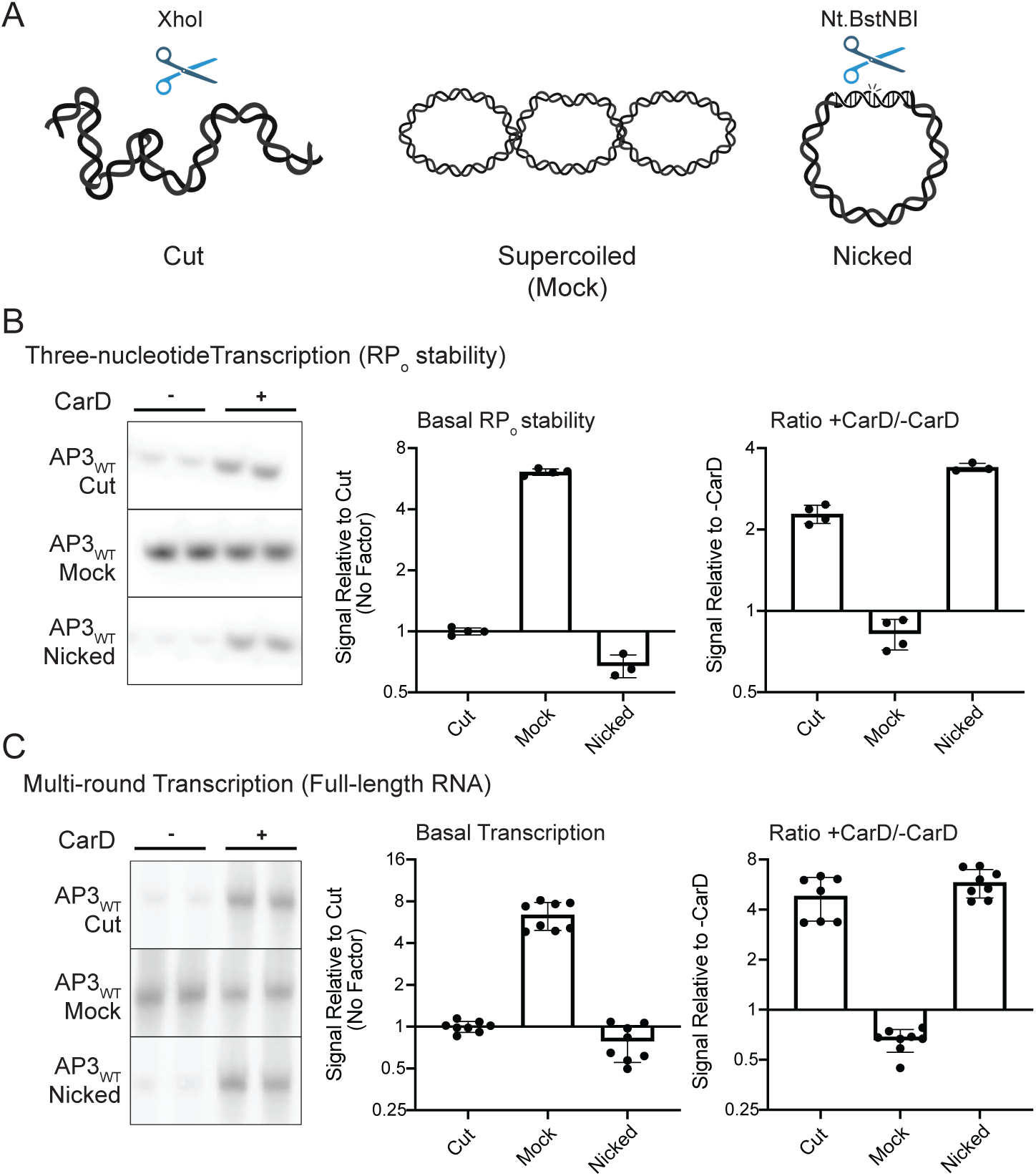
DNA topology affects the RP_o_ stability of promoters and can alter the regulatory outcome of CarD. (A) Schematic of DNA templates used for *in vitro* transcription reactions. All templates transcribe an identical ∼100-nucleotide RNA product from the wild-type *Mtb rrnA*P3 promoter (AP3_WT_). Each DNA template originates from a negatively supercoiled plasmid that was treated with no enzyme (Mock), a nicking endonuclease (Nicked), or a single-cutting restriction endonuclease (Cut). (B) Representative gels showing [^32^P]-labeled three-nucleotide transcription products formed by *Mtb*RNAP-σ^A^ from each DNA template either in the absence or presence of CarD. (C) Representative gels showing full-length [^32^P]-labeled RNA transcripts produced by *Mtb*RNAP-σ^A^ from each DNA template in either the absence or presence of CarD. (B and C) Bar graphs display (left) the mean basal signal intensity relative to the linear “Cut” DNA template ± standard deviation (SD) and (right) the mean ratio of signal intensity +CarD/-CarD for each promoter ± SD. Group means were compared by one-way ANOVA *p*<0.0001. The full results of pairwise comparisons are listed in **Table S4**.

## DISCUSSION

CarD is an essential transcriptional regulator in *Mtb* that affects the expression of over two-thirds of the genome (8) and whose normal function and expression are required for bacterial survival during various stresses and virulence in mice (9, 10, 13, 49). Numerous *in vitro* studies have shown that CarD stabilizes RP_o_ formed by the housekeeping *Mtb*RNAP-σ^A^ holoenzyme (12, 15, 16), leading to the early model that it functions as a general transcription activator. However, a subsequent RNA-seq study of *Mtb* strains encoding mutant alleles of CarD revealed a more complex scenario where CarD appears to differentially activate or repress transcription from different promoters (8). In an effort to understand how CarD could affect gene expression in a promoter specific manner, we now provide experimental evidence for a relationship between RP_o_ stability and the outcome of CarD regulation that results in promoter specific effects of CarD activity. We find that the ratio of transcript production in multi-round transcription reactions +/- CarD shows a roughly linear inverse correlation with increasing RP_o_ stability, with the most stable promoter variants (AP3_EcoExt_, AP3_Stable_, and AP3_Discr5_) being transcriptionally repressed by CarD. CarD’s effect on mycobacterial transcription *in vivo* also reflects the observations from our *in vitro* experiments where promoters predicted to be repressed by CarD are associated with sequence features that correlate with high RP_o_ stability (extended −10 sequence motif, low GC% discriminator region) and promoters predicted to be activated by CarD are associated with an absence of these features. Collectively, these data support our model in which the specific outcome of CarD-mediated RP_o_ stabilization is dependent on the kinetic properties of a given promoter and not on sequence-specific binding, which could explain the observed differential gene expression effects in CarD mutants *in vivo*.

Our study deepens our understanding of mycobacterial transcription regulation and demonstrates how RNAP-binding factors like CarD add complexity to this process. The true relationship between promoter sequence and CarD regulation is likely more nuanced than the data presented in this study. Although the AP3 promoter variants we generated were designed to increase or decrease RP_o_ stability in a stepwise manner (i.e. AP3_Stable_ is a combination of AP3_Discr_ and AP3_ExoEct_; G+C% is titrated one base at a time in AP3_Discr1_-AP3_Discr5_), the effects of each mutation are likely more complex. Minor base substitutions in a promoter sequence can result in large-scale allosteric effects on other RNAP-DNA interactions (50) and kinetic steps (51) outside of RP_o_. Furthermore, our model was built on the idea that CarD uses a single kinetic mechanism, but CarD’s effects on specific transcription initiation rate constants may differ between promoters. For example, CarD contains a conserved tryptophan residue in its DBD (W85) that is positioned to interact with a T at the −12 position of the non-template DNA strand at the upstream fork of the bubble (12), and it has been hypothesized that this sequence-specific interaction acts as a “wedge” to prevent bubble collapse. In theory, on a promoter lacking T_-12_, CarD’s inhibitory effect on *k_collapse_* may be diminished relative to its effects on the rates of bubble opening and promoter escape, producing a unique kinetic mechanism that is biased towards repression. In our RNA-seq dataset, *Mtb* promoters that were predicted to be repressed by CarD were significantly enriched for non-T bases at the −12 position (8), lending some *in vivo* support for the prediction that this DNA sequence context biases CarD towards transcriptional repression.

The specific interaction between CarD W85 and T_-12_ also raises the possibility that specific promoter DNA sequences could influence CarD’s binding preference for partially opened promoter complexes. A recent study showed that CarD has poor binding affinity for RNAP or DNA alone (52), suggesting that CarD may bind to transcription initiation complexes after the initial association between RNAP and DNA. In our ChIP-seq dataset, *M. smegmatis* promoters associated with a primary TSS and containing a σ^A^-like −10 element motif (A_-11_NNNT_-7_) that were within 100bp of a CarD binding site were significantly under-enriched (hypergeometric test *p*=5.66e-06) for promoters lacking a T_-12_ (22%; 93/429) compared to the genome-wide proportion (30%; 765/2511)(**Table S3, Table S5**). These data support the hypothesis that certain DNA sequence-specific interactions may influence but not determine the association between CarD and transcription complexes at specific mycobacterial promoters.

Another region that we did not study but could affect CarD regulation is the initially transcribed sequence downstream of the TSS, which can affect the kinetics of promoter escape and RNAP pausing (53–55). Simplistically, CarD represses transcription from certain promoters by over-stabilizing RP_o_ and decreasing transcript flux by inhibiting promoter escape (7), leading to an accumulation of abortive transcripts (23). However, this model becomes more complicated when considering a branched pathway of transcription initiation (56), where a fraction of RNAP form moribund complexes that never undergo promoter escape. Kinetic studies using a fluorescent reporter showed that CarD increases the fraction of unescaped RNAP complexes (18), and we show that in some contexts CarD can inhibit the synthesis of a three nucleotide product from RP_o_. These data suggest that CarD could affect steps of initial nucleotide incorporation prior to promoter escape and influence the fraction of RNAP complexes undergoing productive versus moribund transcription (55, 57).

We also find that this relationship between RP_o_ stability and CarD is not only influenced by promoter sequence, where promoters with identical DNA sequence can be differentially activated or repressed depending on their supercoiling status (**Fig. 6**). In our experiments, CarD directly activated transcription from the *Mtb* rRNA promoter AP3 on a linear DNA template but repressed transcription from AP3 on a negatively supercoiled template (**Fig. 6**), which is the predominant topological state of DNA in bacterial cells (47). On the surface, this seems to contradict CarD’s role as a positive regulator of rRNA synthesis *in vivo* (13, 20, 25, 58). However, one possible explanation may be that CarD is required to maintain efficient transcription of operons downstream of highly transcribed regions, such as the rRNA operon, when they accumulate positive supercoils due to their high transcriptional activity (47). We propose that CarD may function to overcome the topologically self-limiting nature of rRNA transcription to promote rapid bacterial growth.

Beyond its role in specifically regulating rRNA synthesis, CarD also affects the transcription of hundreds of other *Mtb* genes *in vivo* (8), which could explain CarD’s pleiotropic effects under different stresses. *Mtb* strains with altered CarD activity are also sensitized to various environmental stresses other than nutrient starvation including oxidative stress, genotoxic stress, and antibiotic treatment (9, 10, 13, 49), but it is still unclear what roles CarD plays under these conditions. The impact of topology and promoter context also implies that CarD may elicit different effects on gene expression in different environments. CarD’s ability to interpret the kinetic properties of a promoter add modularity to the mycobacterial transcription response, because while DNA sequence is essentially constant over the lifetime of a bacterial cell its kinetic properties may be dynamic and responsive to environmental stimuli. *In vivo*, the supercoiling state of a promoter is constantly changing in response to the translocation of polymerases, the enzymatic action of topoisomerases, and DNA damage caused by antibiotics or other genotoxic stresses (47, 59). In addition to DNA supercoiling, other environmental factors such as intracellular NTP concentrations (60) and temperature (61) can influence the kinetic properties of a promoter without affecting DNA sequence. During pathogenesis, *Mtb* may encounter these environmental stimuli in various combinations. Furthermore, the expression of CarD is itself highly responsive to environmental signals including nutrient limitation (49) and DNA damage (9). Understanding how transcription factors like CarD interact with these environmental stresses may provide insight into how *Mtb* responds to the host environment and antibiotic treatment, making this an intriguing direction of future study.

Based on the results of this study, we propose that CarD belongs to a growing class of RNAP-binding transcription factors that include DksA/(p)ppGpp (6, 62), TraR and its phage-encoded homologs (63, 64), and the σ-subunit interacting transcription factors including the Actinobacteria-specific protein RbpA (3, 65–68). Like CarD, these factors coordinate broad transcriptional programs in bacteria (5, 14, 66), highlighting the expanded regulatory range of these factors compared to classical transcription factors that are limited to promoters containing a specific binding motif. All of these global transcriptional regulators function by modulating the kinetics of transcription initiation, albeit via different mechanisms. Whereas CarD stabilizes RP_o_, DksA/(p)ppGpp binds RNAP and destabilizes a kinetic intermediate preceding RP_o_, resulting in transcriptional repression at ribosomal RNA promoters that form unstable RP_o_ and transcriptional activation at promoters of amino acid biosynthesis genes that form relatively stable RP_o_ (6, 62, 69–71). Although they exert opposite effects on initiation kinetics, CarD and DksA/(p)ppGpp share the ability to “read” the kinetic properties of a promoter to exert multiple regulatory outcomes on transcription. This study of CarD’s regulatory mechanism demonstrates how kinetic context influences the activity of this class of RNAP-binding transcription factors and reveals another layer in how bacteria coordinate broad gene expression in response to their environment.

## EXPERIMENTAL PROCEDURES

### Bacterial growth and RNA collection

All *M. smegmatis* strains used in this study were derived from mc^2^155 and grown in LB medium supplemented with 0.5% dextrose, 0.5% glycerol, and 0.05% tween-80 at 37 °C. *M. smegmatis* strains expressing CarD^WT^, CarD^R25E^, CarD^K125E^, or CarD^I27W^ were engineered so that the native copy of *carD* is deleted, and the respective CarD allele is expressed from a constitutive P*myc1-tetO* promoter integrated into the genome. The construction of these strains has been previously described (13, 20). For RNA collection, *M. smegmatis* cultures were grown to OD_600_ 0.5-0.9, pelleted, and lysed in TRIzol reagent (Invitrogen) by bead-beating. RNA was isolated by TRIzol-chloroform extraction followed by isopropanol precipitation and finally resuspended in nuclease-free water (Invitrogen).

### RNA sequencing and data analysis

RNA samples were DNase treated using the TURBO DNA-*free* Kit (Invitrogen) and submitted to the Washington University Genome Technology Access Center for paired-end Illumina sequencing (NovaSeq 6000 XP). Ribosomal RNA was depleted prior to sequencing using the Qiagen FastSelect system. Illumina reads were pre-processed using *FastQC* and adapter sequences were removed using *trimmomatic* (72). Sequencing reads were aligned using *HiSat2* (73) to the *M. smegmatis* mc^2^155 reference genome (assembly ASM1500v1) from the *Ensembl* database (74). Reads mapping to annotated protein coding regions were quantified using *featureCounts* (75). Differential expression analysis was performed using *DESeq2* (76). Downstream data analysis and visualization was performed using custom R scripts.

### Protein purification

Plasmids containing the *M. tuberculosis* H37Rv genomic DNA encoding the different *Mtb* RNAP holoenzyme subunits were a gift from Jayanta Mukhopadhyay (Bose Institute, Kolkata, India). *Mtb*RNAP-σ^A^ holoenzyme was purified as previously described (66, 77). Briefly, *Mtb Mtb*RNAP-σ^A^ holoenzyme protein was expressed in *E. coli* BL21 cells containing the plasmids pET-Duet*-rpoB*-*rpoC* (encoding the β and β’ subunits), pAcYc-Duet-*sigA*-*rpoA* (encoding an N-terminal 10xHis-tagged-σ^A^ subunit and ɑ subunits), and pCDF-*rpoZ* (encoding the ω subunit). Holoenzyme protein was isolated from *E. coli* cell lysate by affinity chromatography using a 2x 5mL HisTrap HP Ni^2+^ affinity columns (Cytiva) and further purified by size exclusion chromatography using a Sephacryl S-300 HiPrep column (Cytiva) to select for associated holoenzyme. Purified *Mtb*RNAP-σ^A^ holoenzyme was flash frozen in storage buffer (50% glycerol, 10mM Tris pH 7.9, 200mM NaCl, 0.1 mM EDTA, 1mM MgCl_2_, 20μM ZnCl, and 2mM DTT) and stored at −80 °C. CarD proteins were expressed in BL21 *E. coli* cells using the pET SUMO vector system described previously (66). Purified CarD protein was stored in 20mM Tris pH 7.9, 150mM NaCl, and 1mM beta-mercaptoethanol.

### *In vitro* transcription

Promoter fragments used for *in vitro* transcription were prepared by annealing two complementary single-stranded DNA oligos (IDT) containing the WT or variant AP3 promoter sequence from positions −39 to +4 relative to the transcription start site to create a linear double-stranded DNA fragment that was ligated into the pMSG434 plasmid. Linear DNA templates used for *in vitro* transcription were prepared PCR amplifying a 437bp fragment from the pMSG434 plasmid. Plasmid DNA templates for *in vitro* transcription were constructed by inserting an intrinsic transcription termination sequence (5’-TTTAT-3’) into the pMSG434 plasmid 70bp downstream of the cloned AP3 transcription start site. Negatively supercoiled plasmids were grown in *E. coli* and then isolated using a QIAGEN Plasmid Midi Kit. To generate cut or nicked plasmid templates, plasmid DNA was incubated with XhoI restriction endonuclease (NEB) at 37 °C or Bt.NstNBI nicking endonuclease (NEB) at 55 °C for 1 hour, respectively. All DNA templates were purified by extracting with Buffer-Saturated Phenol pH >7.4 (Invitrogen) followed by isopropanol precipitation before being used in *in vitro* transcription reactions. A full list of the primers used to construct the DNA templates can be found in **Table S6**. Multi-round *in vitro* transcription assays were performed by combining *Mtb*RNAP-σ^A^ holoenzyme, template DNA, and NTPs in a 20μL reaction volume. Multi-round reactions contained final concentrations of 40nM RNAP holoenzyme, 0.8nM DNA template, 0.1mg/mL BSA, 1mM DTT, 400μM GTP, 200μM ATP, 200μM CTP, 200μM UTP, 20μCi/mL [ɑ-^32^P]-UTP (PerkinElmer), 10mM Tris-HCl pH 7.9, 10mM MgCl_2_, and 40mM NaCl. Reactions were initiated with the addition of NTPs and incubated at 37 °C for 1 hour before being terminated with the addition of 20μL ‘stop buffer’ (95% formamide, and <0.1% bromophenol blue and xylene cyanol). Three nucleotide *in vitro* transcription reactions were performed in the same manner, except with final reaction concentrations of 100nM RNAP holoenzyme, 10nM DNA template, 0.1mg/mL BSA, 1mM DTT, 20μM GpU, 10μM UTP, 62.5μCi/mL [ɑ-^32^P]-UTP, 10mM Tris-HCl pH 7.9, 10mM MgCl_2_, and 40mM NaCl. Multi-round and three nucleotide *in vitro* transcription reaction products were separated by gel electrophoresis on denaturing (7M urea) 8% or 22% polyacrylamide gels, respectively, which were vacuum dried and visualized using a phosphorimager screen. Reactions with CarD contained 25:1 molar ratio CarD:RNAP (1μM CarD for the multi-round *in vitro* transcription reactions or 2.5 μM CarD for the three nucleotide transcription reactions) unless otherwise noted. Full gel images can be found in **Figure S5** and **Figure S6**.

## Data Availability

Raw RNA-seq data has been deposited in the GEO repository under accession code GSE222815.

## Supporting Information

This article contains supporting information.

## Acknowledgements

We thank Drake Jensen, Ana Ruiz Manzano, and Eric Galburt for their helpful discussions and for providing the *E. coli* protein expression strain for *Mtb*RNAP-σ^A^ purification. We also thank John Errico, Helen Blaine, and Daved Fremont for their help in purification of RNA polymerase.

## Author Contributions

**Dennis X. Zhu:** Conceptualization, Methodology, Validation, Formal analysis, Investigation, Writing - Original Draft, Writing - Review & Editing, Visualization. **Christina L. Stallings:** Conceptualization, Methodology, Writing - Original Draft, Writing - Review & Editing, Visualization, Resources, Supervision, Project administration, Funding acquisition.

## Funding and additional information

This work was supported by NIH NIGMS grant GM107544 and a Burroughs Wellcome Fund Investigator in the Pathogenesis of Infectious Disease Award awarded to CLS. DXZ is supported by NIH NIAID grant T32A1007172. We also thank the Genome Technology of NCRR or NIH.

