## Supporting Information for "Transcription regulation by CarD in mycobacteria is guided by basal promoter kinetics"

**Figure S1.** Two replicates from our RNA-sequencing (RNA-seq) experiment were removed from downstream analysis following outlier detection.

**Figure S2.** Comparison of transcript expression changes in CarD RNA polymerase interaction domain (RID) mutants in *M. smegmatis* versus *Mtb* (CarD^R25E^ and CarD^R47E^, respectively) relative to strains expressing wild-type CarD (CarD^WT^).

**Figure S3.** *M. smegmatis* strains expressing CarD RID mutants show largely opposite transcriptomic phenotypes, but there is little correlation in the transcriptomic changes between CarD^R25E^ and CarD^K125E^.

**Figure S4.** Promoter sequence characteristics of *Mtb* and *M. smegmatis* promoters classified by their predicted CarD regulatory outcome determined based on RNA sequencing data.

**Figure S5.** Gel images of reaction products from three-nucleotide *in vitro* transcription experiments performed in this study.

**Figure S6.** Gel images of reaction products from multi-round *in vitro* transcription experiments performed in this study.

**Table S1.** Differential expression analysis of *Mycobacterium smegmatis* strains expressing mutant alleles of CarD (R25E, K125E, I27W) compared to a strain expressing WT CarD. Normalization and differential expression analysis was performed using DESeq2. Columns from left to right: gene.id = *M. smegmatis* MC2155 genome annotation gene ID; baseMean = mean normalized counts across all samples; log2FC.r = log2 fold-change expression in CarDR25E relative to CarDWT; padj.r = adjusted p-values for CarDR25E; log2FC.k = log2 fold-change expression in CarDK125E relative to CarD^WT^; padj.k = adjusted p-values for CarD^K125E^; log2FC.i = log2 fold-change expression in CarD^I27W^ relative to CarD^WT^; padj.i = adjusted p-values for CarD^I27W^; PCA.dim1 = variable contribution to the first principal component in principal component analysis (PCA) (**Fig. S1A**); PCA.dim2 = variable contribution to the second principal component in PCA (**Fig. S1A**).

**Table S2**. List of 757 homologous genes from *Mtb* and *M. smegmatis* that were significantly differentially expressed in both the CarD^R47E^ *Mtb* mutant strain and the CarD^R25E^ *M. smegmatis* mutant strain. Homologs were identified by BLAST of amino acid sequence with an e-value cutoff of 0.01

**Table S3.** (A) Genomic coordinates for the centers 1857 CarD binding sites on the *Mycobacterium smegmatis* chromosome identified by chromatin immunoprecipitation sequencing (ChIP-seq) in Landick *et al.* 2014. The list of sites includes the union of all binding regions identified across two biological replicates of *M. smegmatis* cells expressing CarD-HA. The listed regions show the start and end position of each peak in the MC2155 genome. A full description of the ChIP-seq experiment is detailed in Landick *et al.* 2014. (B) Overlap of transcription start sites (TSSs) with CarD binding sites. Only primary TSSs associated with a protein-encoding gene identified in Martini *et al.* 2019 are listed. Each TSS was classified by their 'predicted.card.outcome' based on their differential expression in CarD^R25E^ and CarD^I27W^ relative to CarD^WT^.

**Table S4.** Statistical tests used to analyze in vitro transcription data were performed using GraphPad Prism Version 9.1.1. All transcript production measurements (both three nucleotide and full-length) were first normalized to the mean level of transcript production from linear WT AP3. Mean normalized basal three nucleotide and full-length transcript production levels were compared using a one-way ANOVA. Post-hoc Tukey's tests were performed to compare normalized transcript production levels from different templates. All CarD activation levels were normalized to reactions with no factor within each DNA template. Mean normalized CarD activation levels were compared using a one-way ANOVA. Post-hoc Tukey's tests were performed to compare normalized CarD activation levels from different templates.

**Table S5.** (A) Sequence analysis of promoter features associated with the *Mtb* primary transcription start sites (TSSs) containing a SigA -10 element motif (A-11NNNT-7). TSSs were identified and named based on their genomic position from Cortes *et al.* 2013. (B) Sequence analysis of promoter features associated with the *M. smegmatis* primary transcription start sites (TSSs) containing a SigA -10 element motif (A-11NNNT-7). TSSs were identified and named based on their genomic position from Martini et al. 2019. **'gene.id'**: downstream gene associated with each TSS from the annotated H37Rv genome. **'sigA.category'**: promoters were categorized as either 'perfect' if they contained a T-12 or 'imperfect' if they contained any other base at position -12 of the -10 element. **'ext.category'**: denotes the presence or absence of a T-15G-14N-13 extended -10 motif. **'predicted.card.outcome'**: TSSs were categorized into one of four classes based on their expression pattern in the (A) CarD^R47E^ and CarD^I27W^ *Mtb* mutants or (B) CarD^R25E^ and CarD^I27W^ mutants. **'discriminator.sequence'**: discriminator sequence was defined as the sequence from the 3' end of the -10 element to the +1 transcription start site. NOTE: if multiple -10 element motifs were present in a promoter, then the discriminator sequence is listed as NA. **'discr.GC'**: proportion of G or C bases in the discriminator sequence

**Table S6.** List of oligos used to construct DNA templates used for *in vitro* transcription. Shaded sequences represent restriction enzyme site overhangs used for cloning. Transcription start sites are **bolded**.

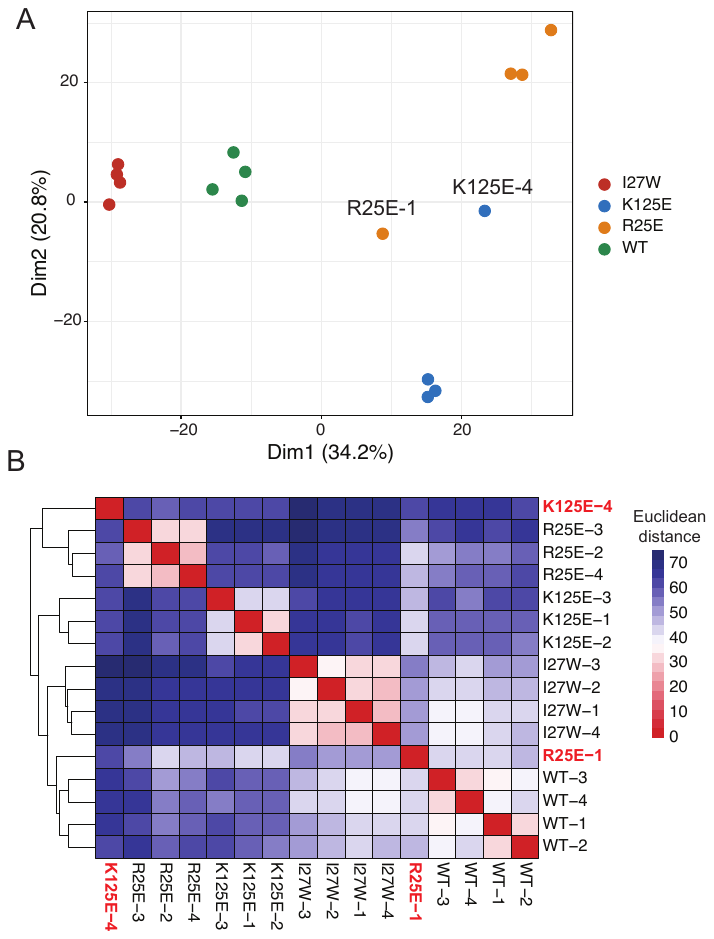

**Figure S1.** Two replicates from our RNA-sequencing (RNA-seq) experiment were removed from downstream analysis following outlier detection. (A) Principal component analysis of RNA-seq samples based on read counts of 6,716 *M. smegmatis* MC^2^155 coding genes. The points are colored by sample genotype. Samples that were removed as outliers are labeled with their sample ID. Both outlier samples were >1.4 standard deviations away from the mean on both of the first two principal components. (B) Clustering of RNA-seq samples based on *M. smegmatis* transcript expression. The heatmap is colored based on the Euclidean distance between pairs of RNA-seq replicates, calculated based on read counts of *M. smegmatis* MC^2^155 coding genes. Outlier samples are colored in red text and cluster with sequencing replicates from different genotypes than their own.

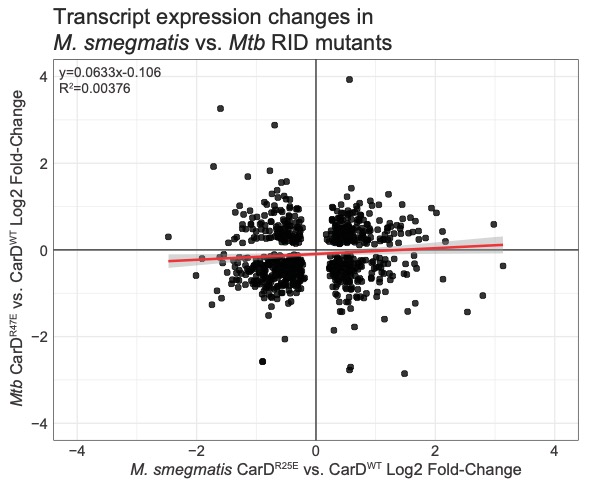

**Figure S2.** Comparison of transcript expression changes in CarD RNA polymerase interaction domain (RID) mutants in *M. smegmatis* versus *Mtb* (CarD^R25E^ and CarD^R47E^, respectively) relative to strains expressing wild-type CarD (CarD^WT^). Points represent 757 homologous protein-encoding genes that were identified by amino acid sequence identity and significantly differentially expressed in both CarD^R25E^ and CarD^R47E^ (**Table S2**). The red line is a linear regression line, and the shaded area around the line represents a 95% confidence interval.

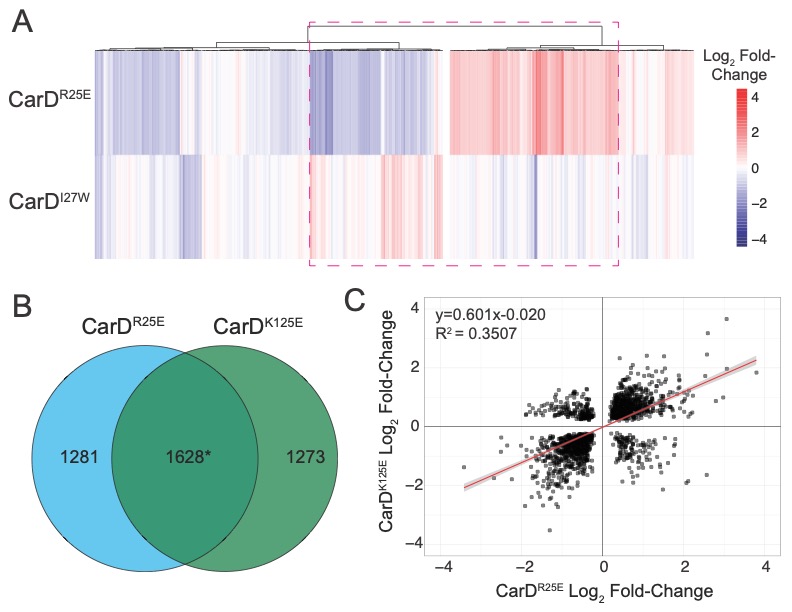

**Figure S3.** *M. smegmatis* strains expressing CarD RID mutants show largely opposite transcriptomic phenotypes, but there is little correlation in the transcriptomic changes between CarD^R25E^ and CarD^K125E^. (A) Hierarchical clustering of 1,186 *M. smegmatis* coding genes that were significantly differentially expressed (*p_adj_* <0.05) greater than 2-fold in at least one CarD mutant genotype. Genes were clustered using Ward’s method based on the log_2_ fold-change in expression in each of the 3 CarD mutant strains relative to CarD^WT^. 158/254 genes differentially expressed greater than 2-fold in CarD^R25E^ are differentially expressed in the opposite direction in CarD^I27W^. The most highly differentially expressed genes in CarD^R25E^ (highlighted by the dashed magenta box) are differentially expressed in an opposite pattern in CarD^I27W^. (B) Venn diagram displaying the overlap in the lists of *M. smegmatis* genes significantly differentially expressed (*p_adj_*<0.05) in CarD^R25E^ versus CarD^K125E^. *The overlap is significant *(p<*0.001) based on a hypergeometric test. (C) Scatter plot displaying transcript expression changes for CarD^K125E^ and CarD^R25E^ on the y- and x-axes, respectively. The red line is a linear regression line, and the shaded area around the line represents a 95% confidence interval.

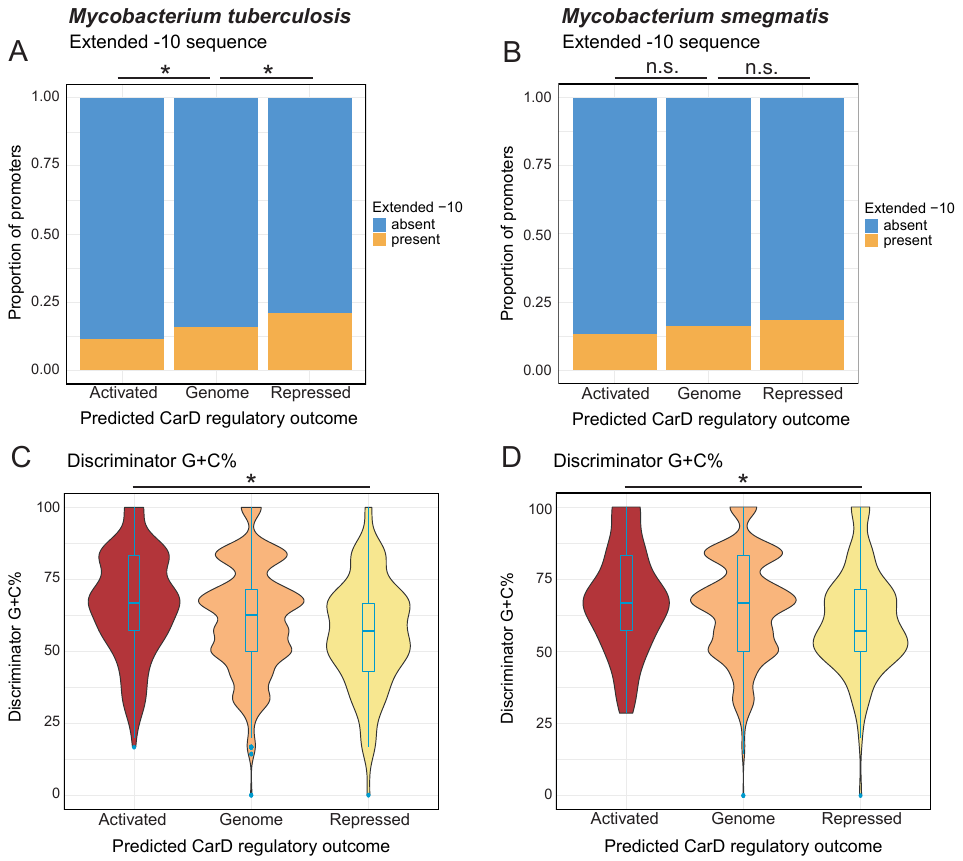

**Figure S4.** Promoter sequence characteristics of *Mtb* and *M. smegmatis* promoters classified by their predicted CarD regulatory outcome determined based on RNA sequencing data. ‘Genome’ denotes the overall proportion of features across all TSSs in the genome. *Mtb* promoters were classified as ‘Activated’ if it was down-regulated in CarD^R47E^ and up-regulated in CarD^I27W^ or ‘Repressed if it was up-regulated in CarD^R47E^ and down-regulated in CarD^I27W^. *M. smegmatis* promoters were classified as ‘Activated’ if it was down-regulated in CarD^R25E^ and up-regulated in CarD^I27W^ or ‘Repressed if it was up-regulated in CarD^R25E^ and down-regulated in CarD^I27W^ (A-B) Bar plots displaying the proportion of promoters containing a consensus extended -10 sequence motif (T_-15_GN_-13_) in either *Mtb* (A) or *M. smegmatis* (B). Enrichment was tested using a hypergeometric test comparing the proportions of each group to the overall proportion of extended -10 promoters among promoters containing a SigA-like -10 sequence motif; * = *p*<0.05, n.s. = not significant. (C-D) Violin plots displaying the discriminator G+C nucleotide base percentage of promoters in *Mtb* (C) or *M. smegmatis* (D). Box and whisker plots displaying the median and interquartile range are drawn in blue. Mean discriminator G+C% values were compared using a Kruskall-Wallis rank sum test followed by post-hoc Dunn’s tests for pairwise comparisons; * = *p*<0.05. The full results of our promoter analysis are listed in **Table S5**.

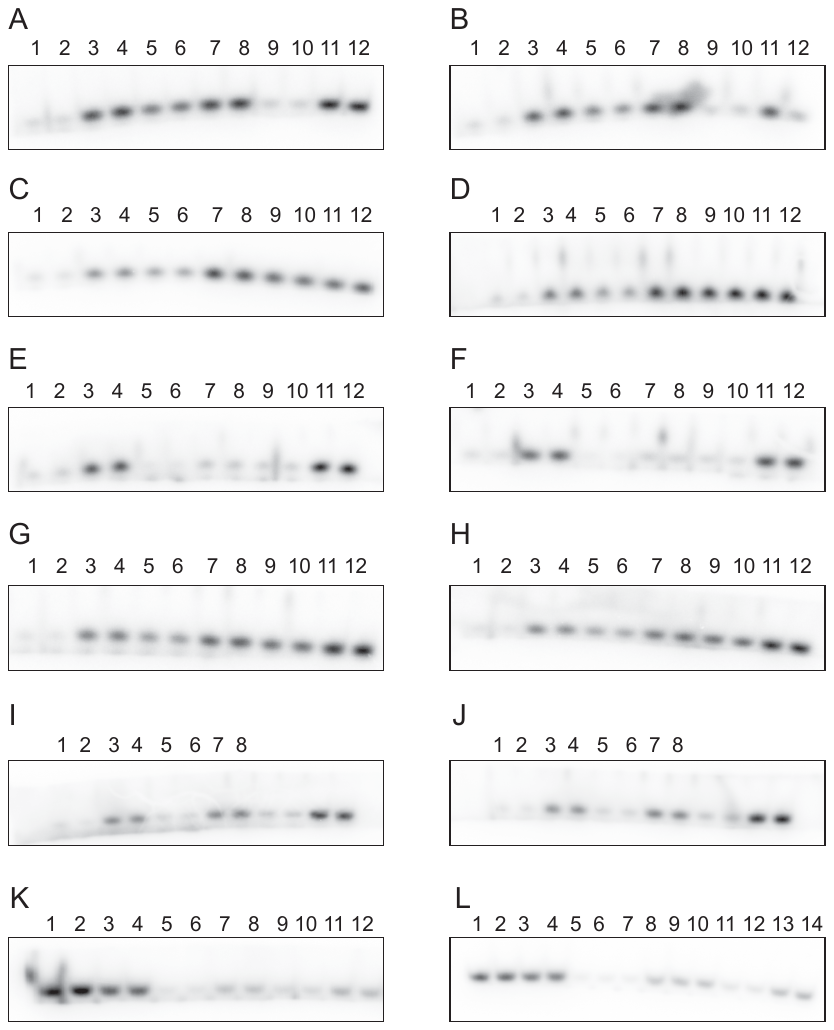

**Figure S5.** Gel images of [^32^P]-labeled three nucleotide *in vitro* transcription reaction products separated on an acrylamide gel and visualized by autoradiography. The samples are AP3_WT_ -CarD (lanes 1-2 on gels A-J), AP3_WT_ +CarD (lanes 3-4 on gels A-J), AP3_MycoExt_ (-CarD A&B 9-10; +CarD A&B 11-12), AP3_Discr_ (-CarD C&D 5-6; +CarD C&D 7-8), AP3_EcoExt_ (-CarD A&B 5-6; +CarD A&B 7-8), AP3_Stable_ (-CarD C&D 9-10; +CarD C&D 11-12), AP3_Discr1_ (-CarD E&F 5-6; +CarD E&F 7-8), AP3_Discr2_ (-CarD E&F 9-10; +CarD E&F 11-12), AP3_Discr3_ (-CarD I&J 5-6; +CarD I&J 7-8), AP3_Discr4_ (-CarD G&H 5-6; +CarD G&H 7-8), AP3_Discr5_ (-CarD G&H 9-10; +CarD G&H 11-12), AP3_Mock_ (-CarD K&L 1-2; +CarD K&L 3-4), AP3_Nicked_ (-CarD K5-6 and L5-7; +CarD K7-8 and L8-10), and AP3_Cut_ (-CarD K9-10 and L11-12; +CarD K11-12 and L13-14). Unnumbered samples in gels I and J represent reactions using promoter templates that were not discussed in this study.

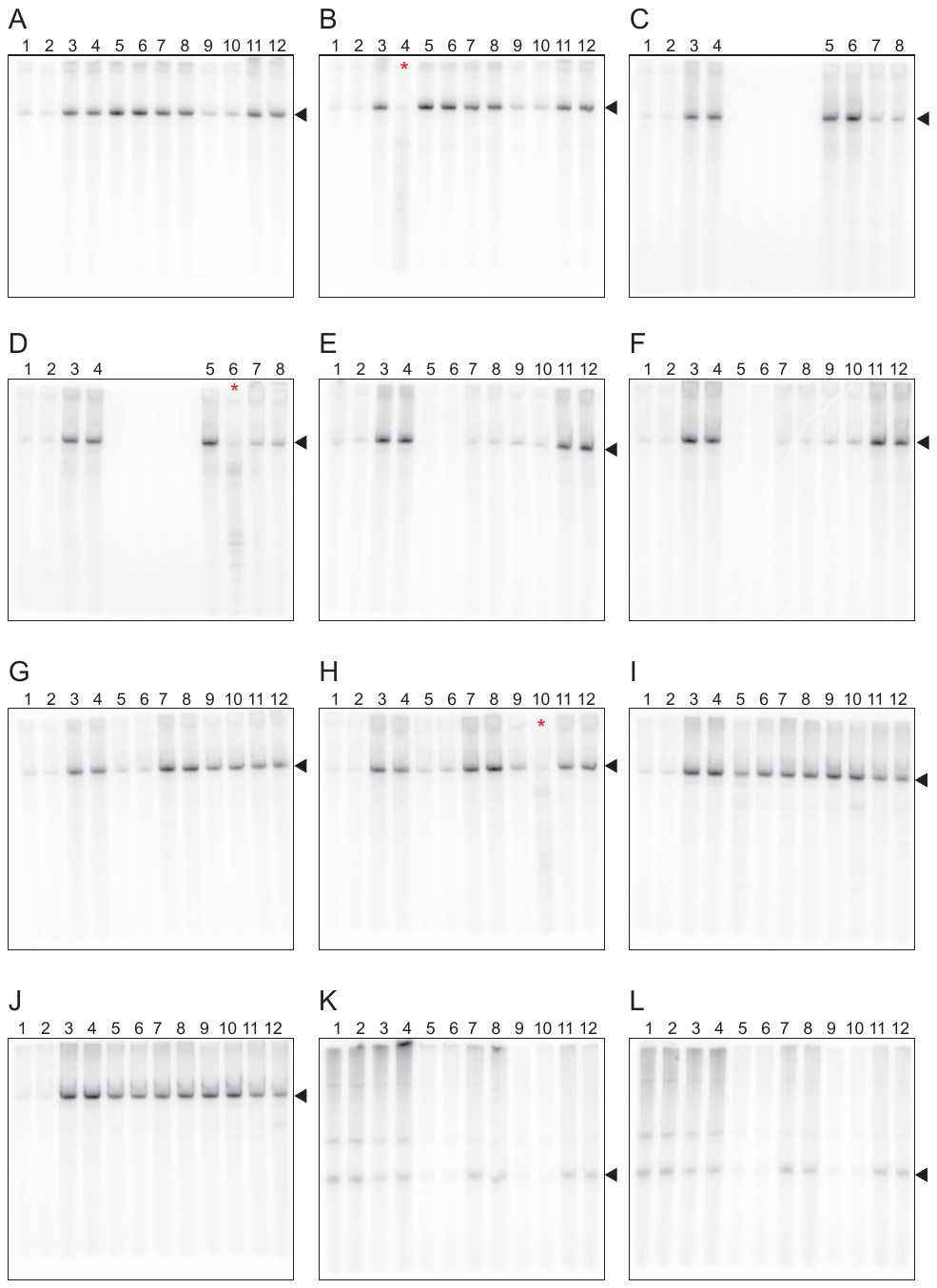

**Figure S6.** Gel images of [^32^P]-labeled multi-round *in vitro* transcription reaction products separated on an acrylamide gel and visualized by autoradiography. The samples are AP3_WT_ -CarD (lanes 1-2 on gels A-J), AP3_WT_ +CarD (lanes 3-4 on gels A-J), AP3_MycoExt_ (-CarD A&B 9-10; +CarD A&B 11-12), AP3_Discr_ (-CarD I&J 5-6; +CarD I&J 7-8), AP3_EcoExt_ (-CarD A&B 5-6; +CarD A&B 7-8), AP3_Stable_ (-CarD C&D 5-6; +CarD C&D 7-8), AP3_Discr1_ (-CarD E&F 5-6; +CarD E&F 7-8), AP3_Discr2_ (-CarD E&F 9-10; +CarD E&F 11-12), AP3_Discr3_ (-CarD G&H 5-6; +CarD G&H 7-8), AP3_Discr4_ (-CarD G&H 9-10; +CarD G&H 11-12), AP3_Discr5_ (-CarD I&J 9-10; +CarD I&J 11-12), AP3_Mock_ (-CarD K&L 1-2; +CarD K&L 3-4), AP3_Nicked_ (-CarD K&L 5-6; +CarD K&L 7-8), and AP3_Cut_ (-CarD K&L 9-10; +CarD K&L 11-12). Black arrows indicate the expected size of promoter-specific products that were used for quantification. Lanes marked with a red asterisk indicate lanes that were affected by RNase degradation and not included in quantification.

| **Name** | **Sequence (5’ to 3’)** |
| --- | --- |
| AP3_WT_ forward primer | GATCCGTCTTGACTCCATTGCCGGATTTGTATTAGACTGGCAGG**G**TTGCA |
| AP3_WT_ reverse primer | TATGCAA**C**CCTGCCAGTCTAATACAAATCCGGCAATGGAGTCAAGACG |
| AP3_EcoExt_ forward primer | GATCCGTCTTGACTCCATTGCCGGATTTGTGTTAGACTGGCAGG**G**TTGCA |
| AP3_EcoExt_ reverse primer | TATGCAACCCTGCCAGTCTAACACAAATCCGGCAATGGAGTCAAGACG |
| AP3_MycoExt_ forward primer | GATCCGTCTTGACTCCATTGCCGGATTTGTAGTAGACTGGCAGG**G**TTGCA |
| AP3_MycoExt_ reverse primer | TATGCAA**C**CCTGCCAGTCTACTACAAATCCGGCAATGGAGTCAAGACG |
| AP3_Discr_ forward primer | GATCCGTCTTGACTCCATTGCCGGATTTGTATTAGACTGGGAGG**G**TTGCA |
| AP3_Discr_ reverse primer | TATGCAA**C**CCTCCCAGTCTAATACAAATCCGGCAATGGAGTCAAGACG |
| AP3_stable_ forward primer | GATCCGTCTTGACTCCATTGCCGGATTGTGTTAGACTGGGAGG**G**TTGCA |
| AP3_stable_ reverse primer | TATGCAA**C**CCTCCCAGTCTAACACAATCCGGCAATGGAGTCAAGACG |
| AP3_Discr1_ forward primer | GATCCGTCTTGACTCCATTGCCGGATTTGTATTAGACTGGCCGG**G**TTGCA |
| AP3_Discr1_ reverse primer | TATGCAA**C**CCGGCCAGTCTAATACAAATCCGGCAATGGAGTCAAGACG |
| AP3_Discr2_ forward primer | GATCCGTCTTGACTCCATTGCCGGATTTGTATTAGACTAGCAGG**G**TTGCA |
| AP3_Discr2_ reverse primer | TATGCAA**C**CCTGCTAGTCTAATACAAATCCGGCAATGGAGTCAAGACG |
| AP3_Discr3_ forward primer | GATCCGTCTTGACTCCATTGCCGGATTTGTATTAGACTATCAGG**G**TTGCA |
| AP3_Discr3_ reverse primer | TATGCAA**C**CCTGATAGTCTAATACAAATCCGGCAATGGAGTCAAGACG |
| AP3_Discr4_ forward primer | GATCCGTCTTGACTCCATTGCCGGATTTGTATTAGACTATTAGG**G**TTGCA |
| AP3_Discr4_ reverse primer | TATGCAA**C**CCTAATAGTCTAATACAAATCCGGCAATGGAGTCAAGACG |
| AP3_Discr5_ forward primer | GATCCGTCTTGACTCCATTGCCGGATTTGTATTAGACTATTAAG**G**TTGCA |
| AP3_Discr5_ reverse primer | TATGCAA**C**CTTAATAGTCTAATACAAATCCGGCAATGGAGTCAAGACG |
| pMSG434 linear forward primer | GGCCATGCCTGTCTCGTTGC |
| pMSG434 linear reverse primer | TCAGCTTGGCGGTCTGGGTG |

**Table S6.** List of oligos used to construct DNA templates used for *in vitro* transcription. Shaded sequences represent restriction enzyme site overhangs used for cloning. Transcription start sites are **bolded**.
